# DUX4-induced HSATII RNA accumulation drives protein aggregation impacting RNA processing pathways

**DOI:** 10.1101/2024.12.17.628988

**Authors:** Tessa Arends, Sean R. Bennett, Stephen J. Tapscott

## Abstract

RNA-driven protein aggregation leads to cellular dysregulation, disrupting normal cellular processes, and contributing to the development of diseases and tumorigenesis. Here, we show that double homeobox 4 (DUX4), an early embryonic transcription factor and causative gene of facioscapulohumeral muscular dystrophy (FSHD), induces the accumulation of stable intranuclear RNAs, including nucleolar RNA and human satellite II (HSATII) RNA. Stable intranuclear RNAs drive protein aggregation in DUX4-expressing muscle cells. Specifically, HSATII RNA sequesters RNA methylation factors. HSATII-YBX1 ribonucleoprotein (RNP) complex formation is mediated by HSATII double-stranded RNA and NSUN2 activity. Aberrant HSATII-RNP complexes affect RNA processing pathways, including RNA splicing. Differential splicing of genes mediated by HSATII-RNP complexes are associated with pathways known to be dysregulated by DUX4 expression. These findings highlight the broader influence of DUX4 on nuclear RNA dynamics and suggest that HSATII RNA could be a critical mediator of RNA processing regulation. Understanding the impact of HSATII-RNP formation on RNA processing provides insight into the molecular mechanisms underlying FSHD.

**SUMMARY:** Arends et al. show that DUX4-induction of stable intranuclear RNA, including pericentromeric human satellite II (HSATII) repeat RNA, leads to nuclear protein aggregation. HSATII ribonucleoprotein complexes impact RNA processing downstream of DUX4 expression.

## INTRODUCTION

RNA plays a major role in the formation of ribonucleoprotein (RNP) complexes and the formation of aberrant RNP complexes has been linked to aging, development and several diseases including facioscapulohumeral muscular dystrophy (FSHD) (Cid-Samper et al., 2018; Naskar et al., 2023; Paxman et al., 2022; Shadle et al., 2019; Tauber et al., 2020). In FSHD, aberrant expression of DUX4 plays a critical role in driving both protein aggregation and RNA accumulation, contributing to disease pathology (Arends et al., 2024; Campbell et al., 2023; Feng et al., 2015; Homma et al., 2015; Homma et al., 2016; Shadle et al., 2019). Determining the mechanism underlying DUX4-dependent protein aggregation is crucial to understanding FSHD pathogenesis. Our work identified human satellite II (HSATII) repeat expression and RNA accumulation as a driver of protein aggregation of nuclear proteins, ADAR1 and eIF4A3, and polycomb repressive complexes (Arends et al., 2024; Shadle et al., 2019). DUX4 induced toxicity of muscle cells is in part mediated by HSATII RNA accumulation (Shadle et al., 2019).

These studies indicate that DUX4-induced nuclear noncoding RNAs might have a role in FSHD pathology, normal development, and perhaps cancers that express DUX4. However, little is known regarding the scope of the RNAs and proteins involved in RNP formation.

Human satellites are a high-copy tandem repeat found at pericentromeric regions (Gosden et al., 1975; Tagarro et al., 1994) and are core pericentromeric components that facilitate interactions with DNA-binding proteins to maintain heterochromatin architecture, ensuring chromatin integrity and genome stability (Bierhoff et al., 2014; Bruckmann et al., 2018; Pezer et al., 2012). Expression of human satellite regions occurs in response to stress, heat shock, in early embryonic and senescent cells (Bai et al., 2016; Miyata et al., 2023; Ninomiya et al., 2020; Ninomiya et al., 2021; Yandim and Karakulah, 2019), and has been correlated with genomic instability in cancer and disease (Arends et al., 2024; Hall et al., 2017; Shadle et al., 2019; Smurova and De Wulf, 2018; Ting et al., 2011). Human satellite regions produce functional noncoding RNA that form RNP complexes by acting as scaffolds, sequestering RNA binding proteins (RBPs) to modulate gene regulation (Hall et al., 2017; Iwata et al., 2024; Kishikawa et al., 2016; Ninomiya et al., 2020; Ninomiya et al., 2021; Shadle et al., 2019).

In this study, we interrogate possible mechanisms contributing to protein aggregation and RNA processing dysregulation in a model system of FSHD. Our work demonstrates that DUX4 expression induces accumulation of stable intranuclear RNAs composed of nucleolar-associated RNA and HSATII RNA. Accumulation of stable intranuclear RNAs correlates with disruption in nucleolar architecture. Additionally, nucleolar-associated RNA and HSATII RNA cause nuclear protein aggregation by sequestering certain RBPs forming RNA-specific RNP complexes. Interestingly, HSATII RNA preferentially sequesters m^6^A and m^5^C RNA methylation factors, and their recruitment is dependent upon HSATII RNA accumulation and RNA methylation activity. Our data indicate that HSATII-RNP complexes impact RNA processing pathways including mRNA splicing. The differential splicing of genes mediated by HSATII-RNP complexes is linked to pathways disrupted by DUX4 expression. These findings suggest that DUX4 has a wider impact on nuclear RNA dynamics and imply that HSATII RNA may play a crucial role in regulating RNA processing downstream of DUX4 expression.

## RESULTS

### DUX4 induces accumulation of stable nuclear RNA aggregates

To model the transient expression of DUX4 which occurs at the 4-cell stage in human embryos (Hendrickson et al., 2017), cancers (Smith et al., 2023) and likely in FSHD muscle cells (Snider et al., 2010), we used the MB135iDUX4 cell line, an immortalized human myoblast cell line with a codon altered doxycycline-inducible DUX4 transgene that has been shown to recapitulate the transcriptional program of the endogenous DUX4 in a synchronized population of cells (Jagannathan et al., 2016). We have previously described that both continuous (24-hour doxycycline treatment) and brief (4-hour doxycycline treatment, “dox-pulse”) DUX4 expression in our MB135iDUX4 (“iDUX4”) cell line recapitulates the FSHD gene signature and transcriptional program in cleavage stage embryos (Jagannathan et al., 2016; Resnick et al., 2019). This system allows us to interrogate the mechanisms downstream of DUX4 that contribute to FSHD pathogenesis.

We have previously reported that DUX4 expression in human myoblast cells leads to robust transcription and nuclear accumulation of HSATII RNA (Arends et al., 2024; Shadle et al., 2019). To determine the stability of HSATII nuclear aggregates over time and whether DUX4 induces other nuclear RNA aggregates, we pulsed iDUX4 cells with doxycycline (dox) for four hours and then incubated cells with an analog of uridine, 5-ethynyluridine (EU, an alkyne-modified nucleoside) which is incorporated into nascent RNA, for 16 hours prior to a washout.

Cells were then fixed and analyzed at 24-hour time points (24-, 48-, 72– and 96-hours) (Fig. 1A). DUX4 expression induced accumulation of intranuclear RNA aggregates (as labeled by EU, “EU-RNA”) that persisted up to 72 hours after DUX4 expression (Fig. 1A). Twenty percent of DUX4-expressing cells had intranuclear EU-RNA aggregates by 24 hours (4 hours post washout of EU-labeling). These EU-RNA aggregates remained present at 48 hours (7% cells positive) and 72 hours (3% cells positive) albeit at decreasing frequencies in the overall population (Fig. 1B).

**Figure 1.**
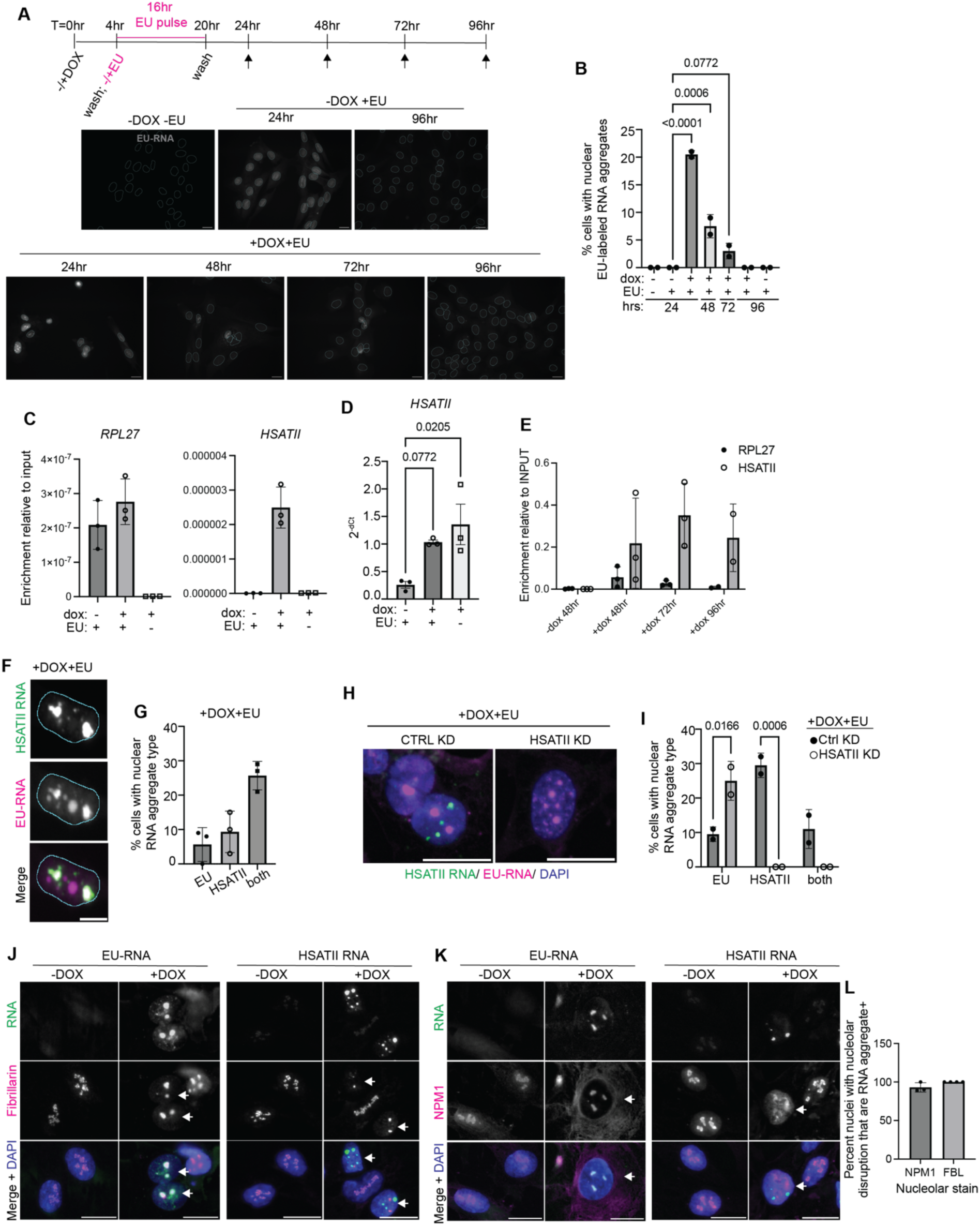
DUX4 induces two types of intranuclear stable RNA species: HSATII RNA and nucleolar-associated RNA. (**A**) Immunofluorescence of EU-RNA aggregates (gray) in iDUX4 cells that were left untreated (-DOX) or pulsed for 4-hours using 1ug/mL doxycycline (+DOX) and then incubated with 0.1mM 5-ethynyluridine (+EU) for 16-hours prior to washout and fixed at 24-hour time points (24-96 hours). Control cells were uninduced (-DOX) iDUX4 cells incubated with EU and fixed at either 24 hours or 96 hours or without EU and fixed at 24 hours. Scale bar = 20μm. Images are representative of two independent experimental replicates performed. (**B**) Percent of cells with intranuclear EU-RNA aggregates in –DOX or +DOX iDUX4 cells labeled with EU for 16-hours at 24-hour time points. EU-RNA aggregates are present within +DOX iDUX4 cells from 24-hour to 72-hour time points. N= 200 nuclei per condition. Data show two independent experimental replicates. Data represent means ± standard deviation (SD). Statistical differences between groups were analyzed employing one-way ANOVA Dunnett’s multiple comparisons test between each group and a control (-DOX+EU 24-hour time point). (**C**) Enrichment of *RPL27* or *HSATII* RNA in isolated EU-RNA from –DOX or +DOX iDUX4 cells incubated with EU for 16-hours or +DOX iDUX4 cells with no EU treatment, all harvested at the 48-hour time point. Housekeeping gene *RPL27* is used as a control. Data show three biological replicates graphed. Data is representative of four independent experimental replicates. Data represent means ± SD. (**D**) Quantitative RT-PCR of *HSATII* expression (2^-ΔCt^) in –DOX or +DOX iDUX4 cells treated with EU for 16-hours or +DOX iDUX4 cells with no EU treatment, all harvested at the 48-hour time point. EU-treatment does not impact *HSATII* RNA expression. Data show three biological replicates graphed. Data is representative of four independent experimental replicates. Data represent means ± SD. Statistical differences between groups were analyzed employing one-way ANOVA Dunnett’s multiple comparisons test between each group and a control (-DOX+EU). (**E**) Enrichment of *RPL27* or *HSATII* RNA in isolated EU-RNA from –DOX or +DOX iDUX4 cells incubated with EU for 16-hours and harvested at 24-hour time points from 48-hours to 96-hours. Housekeeping gene *RPL27* is used as a control. Data show three biological replicates graphed. Data is representative of three independent experimental replicates. Data represent means ± SD. (**F**) Combined immunofluorescence and HSATII RNA-fluorescence in situ hybridization (RNA-FISH) of HSATII RNA (green) and EU-RNA (magenta) in +DOX+EU iDUX4 cells fixed at 48-hour time point. Scale bar = 20μm. Images are representative of three independent experimental replicates performed. (**G**) Percent of cells with intranuclear EU-RNA aggregates only (“EU”), HSATII RNA aggregates only (“HSATII”) or both EU-RNA/ HSATII RNA aggregates (“both”) in +DOX+EU iDUX4 cells fixed at the 48-hour time point. N ≥ 200 nuclei. Dots represent averages of three experimental replicates graphed. Data represent means ± SD. (**H**) Combined immunofluorescence and HSATII RNA-FISH of HSATII RNA (green) and EU-RNA (magenta) in control depleted (“CTRL KD”) or HSATII RNA-depleted (“HSATII KD”) +DOX+EU iDUX4 cells fixed at 48-hour time point. Scale bar = 20μm. Images are representative of two independent experimental replicates performed. (**I**) Percent of cells with intranuclear RNA aggregates: EU only, HSATII only or both in CTRL KD or HSATII KD +DOX+EU iDUX4 cells fixed at the 48-hour time point. N ≥ 200 nuclei. Dots represent averages of two experimental replicates graphed. Data represent means ± SD. Statistical differences between groups were analyzed employing two-way ANOVA Šídák’s multiple comparisons test between CTRL KD and HSATII KD within each group (EU, HSATII, both). (**J**) Combined immunofluorescence of EU-RNA (green) and Fibrillarin (magenta) in –DOX+EU or +DOX+EU iDUX4 cells fixed at 48-hour time point or combined immunofluorescence and HSATII RNA-FISH of HSATII RNA (green) and fibrillarin (magenta) in –DOX and +DOX iDUX4 cells fixed at 24-hours. Scale bar = 20μm. Arrows indicate nuclei with disrupted nucleolar staining. Images are representative of two independent experimental replicates performed. (**K**) Immunofluorescence of EU-RNA (green) and nucleophosmin-1 (NPM1) (magenta) in –DOX+EU or +DOX+EU iDUX4 cells fixed at 48-hour time point or combined immunofluorescence and HSATII RNA-FISH of HSATII RNA (green) and NPM1 (magenta) in –DOX and +DOX iDUX4 cells fixed at 24-hours. Scale bar = 20μm. Arrows indicate nuclei with disrupted nucleolar staining. Images are representative of two independent experimental replicates performed. (**L**) Percent of cells with nucleolar disruption present in cells with intranuclear RNA aggregates. N ≥ 300 nuclei. Dots represent averages of fields analyzed. Two experimental replicates performed.

### Two types of nuclear RNA aggregates induced by DUX4: HSATII RNA aggregates and nucleolar-associated RNA aggregates

We isolated EU-RNA using Click-iT chemistry which captures with high efficiency and sensitivity EU-labeled molecules by a click reaction between a biotin azide and EU terminal alkyne and then subsequently isolated using streptavidin magnetic beads. We isolated EU-RNA at the 48-hour timepoint and performed quantitative RT-PCR (RT-qPCR) for HSATII. HSATII RNA was enriched in isolated EU-labeled RNA from DUX4-expressing cells, compared to RPL27 RNA which had similar enrichment between uninduced (no dox induction) and DUX4-expressing cells (Fig. 1C). To show that EU-labeling did not impact DUX4-induced HSATII RNA expression, RT-qPCR of input RNA showed equivalent HSATII RNA levels in control and EU-labeled DUX4-expressing cells (Fig. 1D). Time course experiments revealed that EU-labeled HSATII RNA remained enriched up to 72 hours post DUX4-expression and started to decline by 96 hours, compared to EU-labeled RPL27 RNA which quickly declined after 48 hours (Fig. 1E).

Microscopy analysis revealed co-localization of HSATII RNA and EU-labeled RNA within DUX4-expressing cells, as well as distinct EU-labeled RNA foci that did not localize with HSATII RNA, which we will refer to as EU-RNA foci to distinguish them from the HSATII-associated EU-labeled foci, now referred to as HSATII RNA foci (Fig. 1F). Many nuclei in dox-pulsed iDUX4 cells contained both EU-RNA and HSATII RNA foci (26%) compared to HSATII RNA foci only (9%) or EU-RNA foci only (6%) (Fig. 1G). This suggested that HSATII RNA was enriched in the EU-RNA population but did not constitute all the stable RNA aggregates. Additionally, depletion of HSATII RNA using antisense oligonucleotides (ASOs) did not impact accumulation of all stable EU-RNA foci, where 10% of control knockdown cells were EU-RNA foci positive compared to 25% HSATII-depleted cells were EU-RNA foci positive (Fig. 1H and I).

The subset of stable EU-RNA foci that did not localize with HSATII RNA specifically co-localized with nucleolar proteins, which we will now refer to as “nucleolar RNA foci.” DUX4-induced nucleolar RNA foci localized with fibrillarin, which is enriched in the dense fibrillar component of the nucleolus, and nucleophosmin-1 (NPM1) which is enriched in the granular component of the nucleolus (Fomproix et al., 1998; Frottin et al., 2019; Lafontaine and Tollervey, 2000) (Fig. 1J and K). HSATII RNA showed no localization with either fibrillarin or NPM1 (Fig. 1J and K). Interestingly, DUX4 expression disrupted the localization of fibrillarin and NPM1, where fibrillarin staining became more punctate with distinct round foci and NPM1 showed disrupted nucleolar staining and increased cytoplasmic signal (Fig. 1J and K). This disruption in nucleolar staining only occurred in nuclei with DUX4-induced intranuclear RNA aggregates (Fig. 1L). The nucleolus is a site for ribosomal RNA (rRNA) biogenesis (Boisvert et al., 2007), thus we next determined whether rRNAs were enriched in the DUX4-induced stable EU-RNA foci. 45S precursor rRNA expression was slightly elevated in DUX4-expressing cells, where 28S mature rRNA were unchanged between conditions (Fig. S1A). Isolated EU-RNA from dox-pulsed iDUX4 cells showed enrichment of 45S precursor rRNA but not mature 28S rRNA (Fig. S1B), similar to the enrichment of HSATII RNAs in the stable EU RNAs (Fig. S1C), suggesting that some rRNAs may constitute a portion of the stable intranuclear EU-RNA foci (Fig. S1B). These data indicate that DUX4 induces accumulation of distinct nuclear RNA foci, either nucleolar RNA foci which comprises 45S RNA, or HSATII RNA foci, that persist for days following DUX4 expression.

### EU-RNA pulldown identifies protein components of DUX4-induced stable ribonucleoprotein complexes

To determine the protein composition of DUX4-induced stable intranuclear ribonucleoprotein (RNP) complexes, we modified the RNA interactome using click chemistry (RICK) approach (Bao et al., 2018). To capture DUX4-induced stable EU-labeled RNA associated RNP complexes (EU-RNPs), iDUX4 cells were either pulsed with dox for 4 hours or not treated and then incubated with EU for 16 hours prior to a second washout and terminally harvested at 48 hours. Cells were UV crosslinked to preserve interactions between RNA and proteins and EU-RNA and any associated RNP complexes were isolated, and the associated protein was subjected to mass spectrometry.

Mass spectrometry of EU-RNA associated proteins identified 105 proteins enriched in the DUX4-induced stable EU-RNPs (>2 unique peptide matches and >1.5 fold difference; Suppl. Table 1). Gene set enrichment using Enrichr (Chen et al., 2013; Kuleshov et al., 2016; Xie et al., 2021) found that the DUX4-induced stable EU-RNA associated proteins were enriched in biological processes including mRNA splicing, mRNA processing, RNA stabilization, regulation of gene expression and regulation of translation (Fig. 2A). Some of these proteins were previously shown to have disrupted localization and induced aggregation in DUX4-expressing muscle cells, including TDP-43 and SRSF2/SC35 (Homma et al., 2015; Homma et al., 2016), and we validated their enrichment in the dox-treated EU-RNA fraction by immunoblot (Fig. 2B). Of particular interest, several factors involved in 5-methylcytosine (m^5^C) RNA regulation including RNA methyltransferase NSUN2 and m^5^C-reader YBX-1 (Chen et al., 2019; Hussain et al., 2013; Yang et al., 2017; Yang et al., 2019) were also enriched in these RNPs and validated by immunoblot (Fig. 2B).

**Figure 2.**
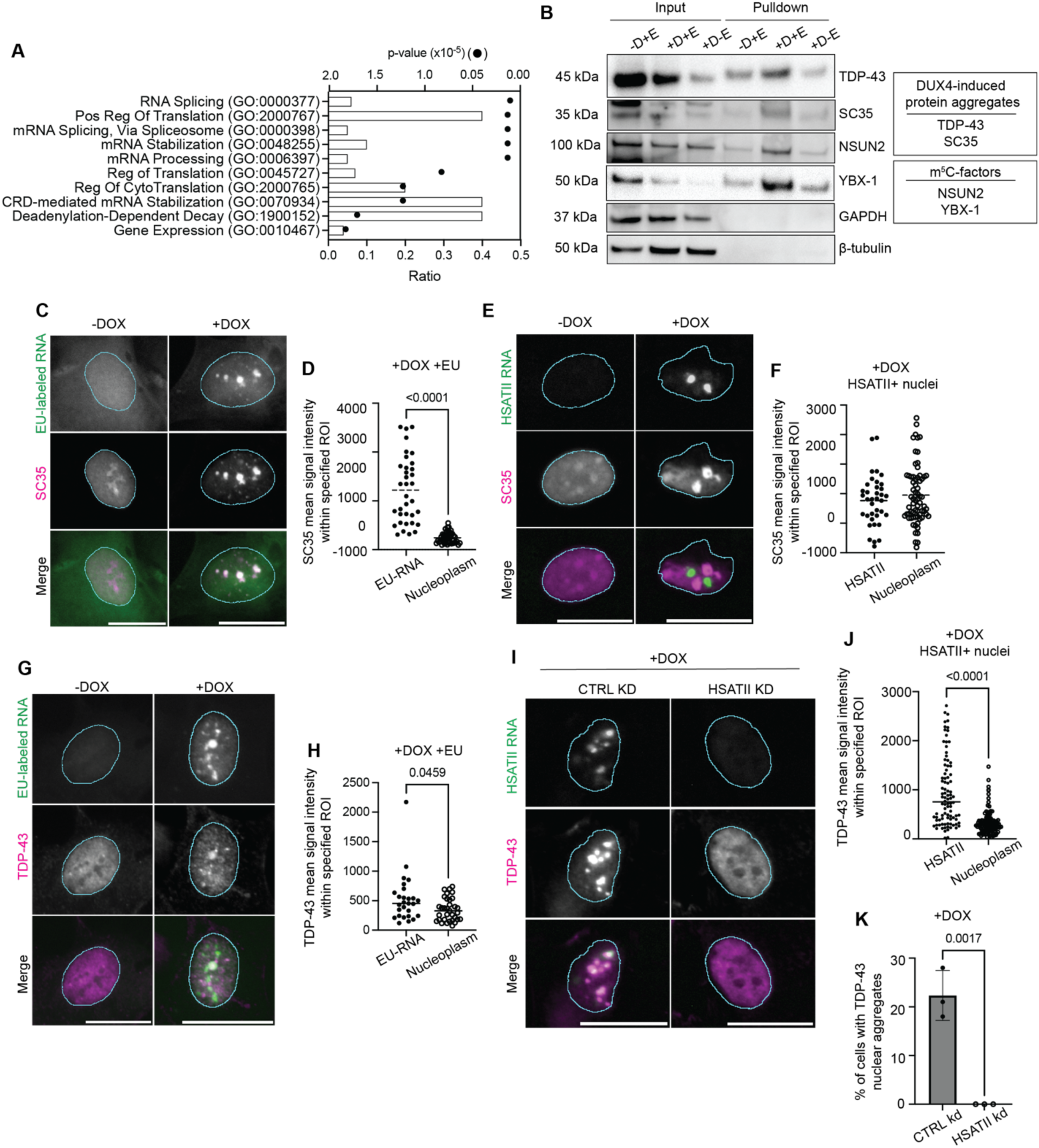
Distinct ribonucleoprotein complexes are formed with nucleolar-associated EU-RNA and HSATII RNA aggregates. (**A**) Biological process pathway analysis of DUX4-induced stable EU-RNA associated RNPs (EU-RNPs) proteomics filtered protein hits (>2 unique peptide matches and >1.5 difference) ran through Enrichr. EU-RNP complexes were isolated from – DOX+EU, +DOX+EU or +DOX-EU iDUX4 cells harvested at the 48-hour time point using the “RICK” approach (Bao et al., 2018). Protein was purified from isolated EU-RNP complexes and was subjected to mass-spectrometry. Proteomics was performed in experimental duplicates. (**B**) Validation of selected proteins identified through EU-RNP proteomics: Known DUX4-induced protein aggregates SC35 and TDP-43, and m^5^C-related factors NSUN2 and YBX-1. Proteins were isolated as stated in Fig. 2A. GAPDH and beta-tubulin were used as loading controls for inputs and negative controls for pulldown samples. Blot shows singletons, but representative of experimental triplicates. –D+E, –DOX+EU samples; +D+E, +DOX+EU samples; +D-E, +DOX-EU samples. (**C**) Immunofluorescence of EU-RNA (green) and SC35 (magenta) in –DOX+EU or +DOX+EU iDUX4 cells fixed at 48-hour time point. Scale bar = 20μm. Images are representative of two independent experimental replicates performed. (**D**) Mean signal intensity of SC35 within specified regions of interest (ROI) in the nucleus: either within EU-RNA foci or the remainder of the nucleoplasm in +DOX EU-RNA+ nuclei in iDUX4 cells. Dots are each individual ROI. N ≥ 35 ROI. ROI within the nucleoplasm is drawn with relatively the same circumference as EU-RNA foci ROI. Data are representative of two independent experimental replicates. Data represent means. Statistical differences between groups were analyzed employing Mann Whitney test. (**E**) Combined immunofluorescence and HSATII RNA-FISH of HSATII (green) and SC35 (magenta) in –DOX or +DOX iDUX4 cells fixed at 24-hour time point. Scale bar = 20μm. Images are representative of two independent experimental replicates performed. (**F**) Mean signal intensity of SC35 within specified ROI in the nucleus: either within HSATII RNA foci or the remainder of the nucleoplasm in +DOX HSATII+ nuclei in iDUX4 cells. Dots are each individual ROI. N ≥ 35 ROI. ROI within the nucleoplasm is drawn with relatively the same circumference as HSATII RNA foci ROI. Data are representative of two independent experimental replicates. Data represent means. (**G**) Immunofluorescence of EU-RNA (green) and TDP-43 (magenta) in –DOX+EU or +DOX+EU iDUX4 cells fixed at 48-hour time point. Scale bar = 20μm. Images are representative of two independent experimental replicates performed. (**H**) Mean signal intensity of TDP-43 within specified ROI in the nucleus: either within EU-RNA foci or the remainder of the nucleoplasm in +DOX EU-RNA+ nuclei in iDUX4 cells. Dots are each individual ROI. N ≥ 30 ROI. ROI within the nucleoplasm is drawn with relatively the same circumference as EU-RNA foci ROI. Data are representative of two independent experimental replicates. Data represent means. Statistical differences between groups were analyzed employing Mann Whitney test. (**I**) Combined immunofluorescence and HSATII RNA-FISH of HSATII (green) and TDP-43 (magenta) in –DOX or +DOX iDUX4 cells with CTRL KD or HSATII KD fixed at 24-hour time point. Scale bar = 20μm. Images are representative of three independent experimental replicates performed. (**J**) Mean signal intensity of TDP-43 within specified ROI in the nucleus: either within HSATII RNA foci or the remainder of the nucleoplasm in +DOX HSATII+ nuclei in iDUX4 cells. Dots are each individual ROI. N ≥ 90 ROI. ROI within the nucleoplasm is drawn with relatively the same circumference as HSATII RNA foci ROI. Data are representative of three independent experimental replicates. Data represent means. Statistical differences between groups were analyzed employing Mann Whitney test. (**K**) Percent of cells with TDP-43 nuclear aggregates in +DOX iDUX4 cells with CTRL KD or HSATII KD. N ≥ 200 nuclei. Dots represent averages of three experimental replicates graphed. Data represent means ± SD. Statistical differences between groups were analyzed employing two-tailed Unpaired t-test.

To determine whether identified RNA binding proteins were associated with either nucleolar RNA foci or HSATII RNA foci, we determined their localization using microscopy. Microscopy analysis revealed that SC35 specifically co-localized with nucleolar RNA foci (Fig. 2C). SC35 nuclear signal was significantly enriched within EU+ nucleolar RNA foci (mean signal intensity: 1120 ± 685) compared with random regions of interest (ROI) within the nucleoplasm of EU-RNA+ nuclei (243 ± 111) (Fig. 2D). SC35 did not localize with HSATII RNA (Fig. 2E), where SC35 signal intensity was not significantly changed between HSATII RNA foci (881 ± 422) and ROI within the nucleoplasm in HSATII RNA+ nuclei (974 ± 494) (Fig. 2F). In contrast, TDP-43 did not localize with all EU+ nucleolar RNA foci, where TDP-43 nuclear signal intensity was similar between EU+ nucleolar RNA foci (528 ± 408) and ROI within the nucleoplasm of EU-RNA+ nuclei (349 ± 195) (Fig. 2G and H); whereas TDP-43 strongly co-localized with HSATII RNA foci (Fig. 2I). TDP-43 signal intensity was significantly enriched in HSATII RNA foci (949 ± 700) compared to ROI in the nucleoplasm of HSATII RNA+ nuclei (327 ± 250) (Fig. 2J).

To determine whether HSATII RNA accumulation was necessary for TDP-43 nuclear aggregation, we depleted cells of HSATII RNA using ASOs. Depletion of HSATII RNA completely diminished nuclear aggregation of TDP-43 (Fig. 2I), where 20% of control ASO treated cells contained TDP-43 nuclear aggregates compared to HSATII-depleted cells where no TDP-43 aggregates were observed (Fig. 2K). Therefore, the nuclear aggregation of SC35 and TDP-43 that was previously observed in FSHD muscle cells is due to sequestration by nucleolar RNA or HSATII RNA, respectively.

### HSATII RNA associates with m5C– and m6A-related RNA methylation factors

To specifically identify the HSATII RNA-protein interactome, as opposed to all stable EU-labeled RNAs, we employed a modified version of chromatin isolation by RNA purification (ChIRP) approach and used biotinylated oligonucleotides complementary to HSATII sequences to pull-down endogenous HSATII-RNP complexes (Fig. 3A) (Chu et al., 2012). We either doxycycline-treated iDUX4 cells or left iDUX4 cells untreated and harvested 20 hours after a 4-hour doxycycline pulse. We used our previously validated HSATII RNA probes which efficiently capture all HSATII RNA (Arends et al., 2024; Shadle et al., 2019) or used control probes. Compared to pull-down with a control ASO, RT–qPCR analysis demonstrated robust enrichment of HSATII RNA with the HSATII ASO from dox-pulsed iDUX4 cells whereas neither *ZSCAN4*, a DUX4-induced gene, or *RPL27* mRNA was isolated with the HSATII-ASO (Fig. 3B). We also verified protein enrichment of known HSATII RNA interactors eIF4A3 and MeCP2 (Fig. 3C) (Hall et al., 2017; Shadle et al., 2019). Proteomics identified over 300 proteins (>10 unique peptide matches within +DOX HSATII ASO and >1.5 fold difference between +DOX HSATII ASO and control ASO) associated with HSATII RNA from dox-pulsed iDUX4 cells, most of which were not previously reported as HSATII interacting factors (Suppl. Table 2). Gene ontology analysis using Enrichr indicated significant enrichment of proteins involved in ribosome biogenesis, ribonucleoprotein complex biogenesis, RNA splicing, RNA modification and noncoding RNA processing (Fig. 3D).

**Figure 3.**
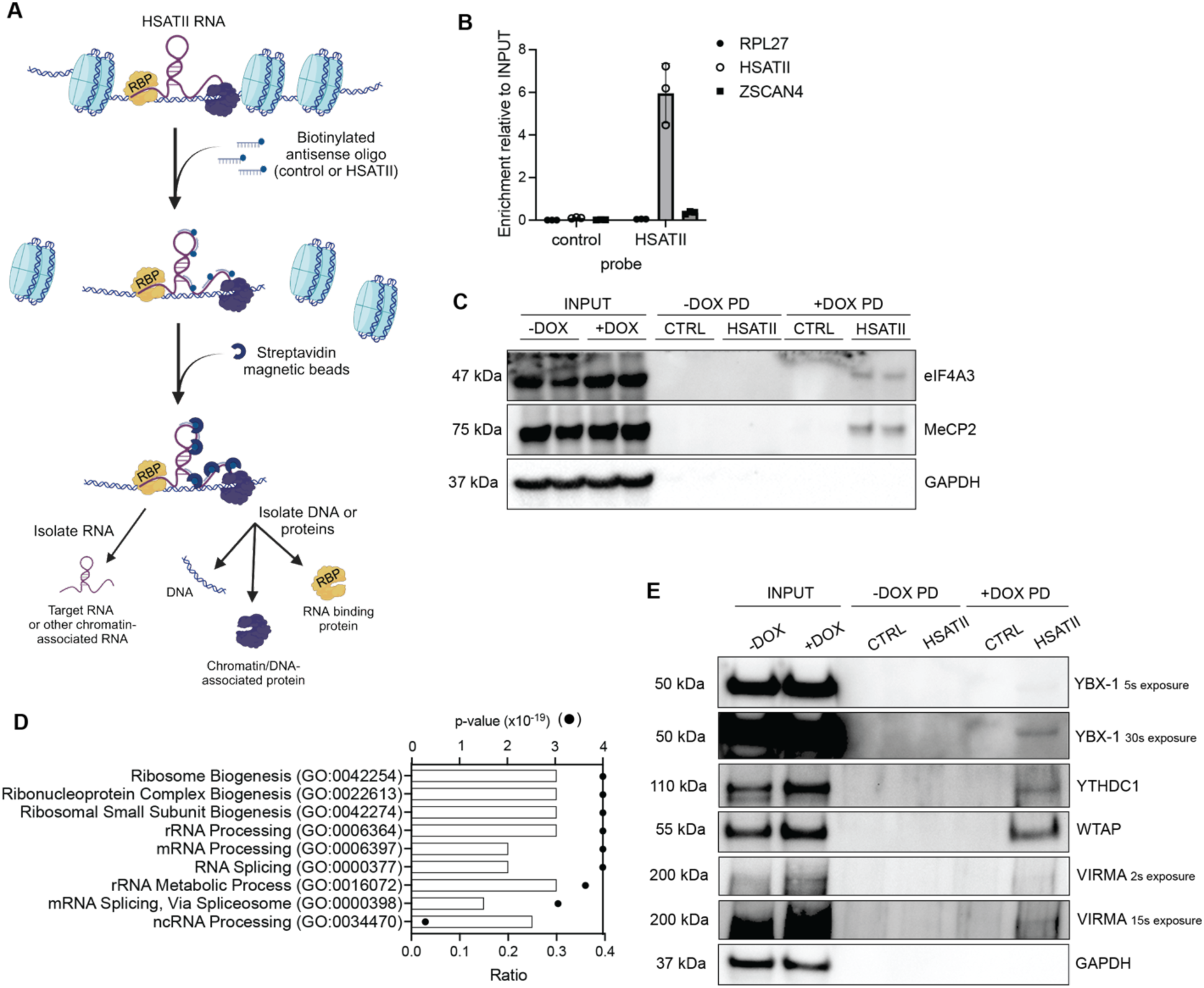
Identification of HSATII RNA interacting proteins. (**A**) Experimental schematic showing ChIRP protocol (Chu et al., 2012). (**B**) RT-qPCR showing enrichment of *RPL27*, *ZSCAN4* or *HSATII* RNA in ChIRP pulldowns using either control ASOs or HSATII-specific ASOs in +DOX iDUX4 cells. N=3 experimental replicates. (**C**) Immunoblot showing enrichment of known HSATII interacting proteins MeCP2 and eIF4A3 in ChIRP pulldowns (PD) using either control ASOs (CTRL) or HSATII-specific ASOs (HSATII) in –DOX or +DOX iDUX4 cells. GAPDH used as loading control for inputs and negative control for PD samples. Immunoblot shows two biological replicates. N=3 experimental replicates performed. (**D**) Biological process pathway analysis of HSATII RNA associated RNPs (HSATII-RNP) proteomics filtered protein hits (>2 unique peptide matches and >1.5 difference) ran through Enrichr. HSATII-RNP complexes were isolated from –DOX or +DOX iDUX4 cells harvested at the 24-hour time point using the “ChIRP” approach (Chu et al., 2012). Protein was purified from isolated HSATII-RNP complexes using antisense oligonucleotides (ASO) targeting HSATII or control sequences and was subjected to mass-spectrometry. Proteomics was performed in biological triplicate. (**E**) Validation of selected proteins identified through HSATII-RNP proteomics: m^5^C-related factors YBX-1 and m^6^A-related factors YTHDC1, WTAP, and VIRMA. Proteins were isolated as stated in Fig. 3B. GAPDH was used as a loading control for inputs and negative control for pulldown samples. Blot shows singletons, but representative of experimental triplicates. CTRL, Control ASO; HSATII, HSATII-specific ASO; PD, pulldown.

Several known HSATII interacting proteins were identified in our mass spectrometry analysis, including MeCP2, eIF4A3 and ADAR1 (Suppl. Table 2) (Hall et al., 2017; Shadle et al., 2019). Notably, RNA methylation related factors were identified, including factors involved in m^5^C (NSUN2 and YBX-1) and factors involved in N^6^-methyladenosine (m^6^A) (VIRMA, WTAP and YTHDC1) (Suppl. Table 2) (Chellamuthu and Gray, 2020; Hussain et al., 2013; Yang et al., 2017; Zaccara et al., 2019). Verification of identified HSATII-interacting proteins was confirmed by immunoblotting of ChIRP-enriched proteins (Fig. 3E). These findings underscore selective interactions of RNA binding proteins, particularly of RNA methylation-related proteins, with HSATII RNA.

### m^6^A-related factors are sequestered by HSATII RNA

Interestingly, m^6^A-related factors have been shown to interact with human satellite III RNA in cells undergoing stress response and recovery (Ninomiya et al., 2021; Timcheva et al., 2022), and major satellite transcripts were found to be enriched for m^6^A-modification in mouse embryonic stem cells (Duda et al., 2021). This suggests a possible conserved function of satellite RNA to associate with m^6^A-related factors. To further validate the association and localization of HSATII RNA with identified m^6^A-factors, including Vir-like m^6^A methyltransferase associated (KIAA1429/VIRMA), Wilms tumor 1-associated protein (WTAP), and YTH domain-containing protein 1 (YTHDC1), we performed immunofluorescence staining combined with HSATII RNA-fluorescence *in situ* hybridization (RNA-FISH) (Fig. 4). m^6^A writer complex component VIRMA, showed strong co-localization with HSATII RNA foci in dox-pulsed iDUX4 cells (Fig. 4A), where 74% of HSATII+ nuclei had VIRMA nuclear aggregates that at least partially overlapped with HSATII RNA foci (Fig. 4B). VIRMA nuclear signal was specifically enriched within HSATII RNA foci (1509 ± 458) compared to ROI within the nucleoplasm of HSATII+ dox-pulsed iDUX4 cells (768 ± 454) (Fig. 4C). Nuclear aggregation of VIRMA was dependent on HSATII RNA accumulation because depletion of HSATII RNA using ASOs in dox-pulsed iDUX4 cells abolished VIRMA nuclear aggregates (Fig. 4A); where only 6% of HSATII RNA-depleted cells contained VIRMA nuclear aggregates compared to 23% of control depleted cells (Fig. 4D).

**Figure 4.**
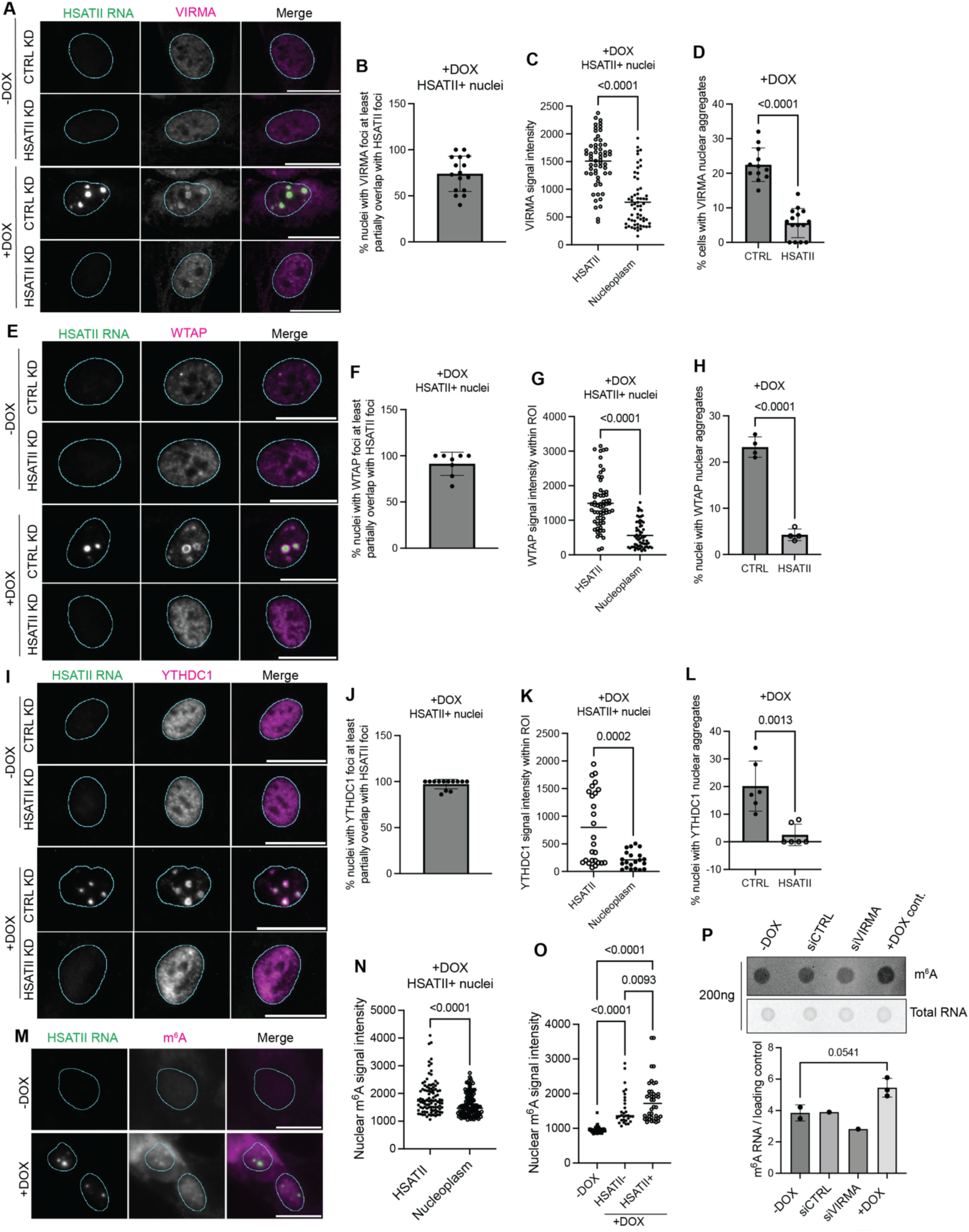
HSATII sequesters and causes nuclear aggregation of m^6^A-related factors. (**A**) Combined immunofluorescence and HSATII RNA-FISH of HSATII (green) and VIRMA (magenta) in –DOX or +DOX iDUX4 cells with CTRL KD or HSATII KD fixed at 24-hour time point. Scale bar = 20μm. Images are representative of three independent experimental replicates performed. (**B**) Percent of nuclei with VIRMA signal that at least have partial signal overlap with HSATII RNA foci in +DOX HSATII+ nuclei in iDUX4 cells. N ≥ 200 nuclei. Dots represent fields taken from representative experiment. Data represent means ± SD. (**C**) Mean signal intensity of VIRMA within specified ROI in the nucleus: either within HSATII RNA foci or the remainder of the nucleoplasm in +DOX HSATII+ nuclei in iDUX4 cells. Dots are each individual ROI. N ≥ 60 ROI. ROI within the nucleoplasm is drawn with relatively the same circumference as HSATII RNA foci ROI. Data are representative of three independent experimental replicates. Data represent means. Statistical differences between groups were analyzed employing Mann Whitney test. (**D**) Percent of cells with VIRMA nuclear aggregates in +DOX iDUX4 cells with CTRL KD or HSATII KD. N ≥ 200 nuclei. Dots represent fields taken from representative experiment. Data represent means ± SD. Data are representative of three independent experimental replicates. Statistical differences between groups were analyzed employing Mann Whitney test. (**E**) Combined immunofluorescence and HSATII RNA-FISH of HSATII (green) and WTAP (magenta) in –DOX or +DOX iDUX4 cells with CTRL KD or HSATII KD fixed at 24-hour time point. Scale bar = 20μm. Images are representative of three independent experimental replicates performed. (**F**) Percent of nuclei with WTAP signal that at least have partial signal overlap with HSATII RNA foci in +DOX HSATII+ nuclei in iDUX4 cells. N ≥ 200 nuclei. Dots represent fields taken from representative experiment. Data represent means ± SD. (**G**) Mean signal intensity of WTAP within specified ROI in the nucleus: either within HSATII RNA foci or the remainder of the nucleoplasm in +DOX HSATII+ nuclei in iDUX4 cells. Dots are each individual ROI. N ≥ 60 ROI. ROI within the nucleoplasm is drawn with relatively the same circumference as HSATII RNA foci ROI. Data are representative of three independent experimental replicates. Data represent means. Statistical differences between groups were analyzed employing Mann Whitney test. (**H**) Percent of cells with WTAP nuclear aggregates in +DOX iDUX4 cells with CTRL KD or HSATII KD. N ≥ 200 nuclei. Dots represent fields taken from representative experiment. Data represent means ± SD. Data are representative of three independent experimental replicates. Statistical differences between groups were analyzed employing Unpaired t-test. (**I**) Combined immunofluorescence and HSATII RNA-FISH of HSATII (green) and YTHDC1 (magenta) in –DOX or +DOX iDUX4 cells with CTRL KD or HSATII KD fixed at 24-hour time point. Scale bar = 20μm. Images are representative of three independent experimental replicates performed. (**J**) Percent of nuclei with YTHDC1 signal that at least have partial signal overlap with HSATII RNA foci in +DOX HSATII+ nuclei in iDUX4 cells. N ≥ 200 nuclei. Dots represent fields taken from representative experiment. Data represent means ± SD. (**K**) Mean signal intensity of YTHDC1 within specified ROI in the nucleus: either within HSATII RNA foci or the remainder of the nucleoplasm in +DOX HSATII+ nuclei in iDUX4 cells. Dots are each individual ROI. N ≥ 25 ROI. ROI within the nucleoplasm is drawn with relatively the same circumference as HSATII RNA foci ROI. Data are representative of three independent experimental replicates. Data represent means. Statistical differences between groups were analyzed employing Unpaired t-test. (**L**) Percent of cells with YTHDC1 nuclear aggregates in +DOX iDUX4 cells with CTRL KD or HSATII KD. N ≥ 200 nuclei. Dots represent fields taken from representative experiment. Data represent means ± SD. Data are representative of three independent experimental replicates. Statistical differences between groups were analyzed employing Unpaired t-test. (**M**) Combined immunofluorescence and HSATII RNA-FISH of HSATII (green) and m^6^A (magenta) in –DOX or +DOX iDUX4 cells fixed at 24-hour time point. Scale bar = 20μm. Images are representative of three independent experimental replicates performed. (**N**) Mean signal intensity of m^6^A within specified ROI in the nucleus: either within HSATII RNA foci or the remainder of the nucleoplasm in +DOX HSATII+ nuclei in iDUX4 cells. Dots are each individual ROI. N ≥ 100 ROI. ROI within the nucleoplasm is drawn with relatively the same circumference as HSATII RNA foci ROI. Data are representative of three independent experimental replicates. Data represent means. Statistical differences between groups were analyzed employing Mann-Whitney test. (**O**) Mean signal intensity of nuclear m^6^A signal in –DOX or +DOX cells fixed at 24-hour time point. Dots are individual nuclei. N ≥ 50 nuclei. Data are representative of three independent experimental replicates. Data represent means. Statistical differences between groups were analyzed employing One-way ANOVA Tukey’s multiple comparison test. (**P**) RNA dot blot of RNA isolated from uninduced (-DOX), control depleted (siCTRL), VIRMA depleted (siVIRMA) or +DOX iDUX4 cells harvested at 24-hours. 200ng of RNA was loaded. RNA was probed using anti-m^6^A antibody or total RNA was determined using Methylene Blue. Signal intensity showed an increase in m^6^A RNA levels in +DOX iDUX4 cells compared to uninduced cells. Data are representative of three independent experimental replicates.

In addition, the regulatory subunit of the m^6^A writer complex, WTAP, showed co-localization with HSATII RNA foci in dox-pulsed iDUX4 cells (Fig. 4E), where 91% of HSATII+ nuclei had WTAP nuclear aggregates that at least partially overlapped with HSATII RNA foci (Fig. 4F). Within dox-pulsed iDUX4 cells, WTAP nuclear signal was specifically enriched within HSATII RNA foci (1492 ± 722) compared to ROI within the nucleoplasm (563 ± 376) (Fig. 4G). Nuclear aggregation of WTAP was also dependent on HSATII RNA accumulation because depletion of HSATII RNA using ASOs in dox-pulsed iDUX4 cells nearly abolished WTAP nuclear aggregates (Fig. 4E); where only 4% of HSATII RNA-depleted cells contained WTAP nuclear aggregates compared to 23% of control depleted cells (Fig. 4H).

Next, we validated the localization and association of m^6^A-reader YTHDC1 with HSATII RNA. YTHDC1 showed strong co-localization with HSATII RNA foci in dox-pulsed iDUX4 cells (Fig. 4I), where 97% of HSATII+ nuclei had YTHDC1 nuclear aggregate signal overlap with HSATII RNA foci (Fig. 4J). YTHDC1 nuclear signal was specifically enriched within HSATII RNA foci (799 ± 660) compared to ROI within the nucleoplasm of dox-pulsed iDUX4 cells (211 ± 155) (Fig. 4K). Nuclear aggregation of YTHDC1 was dependent on HSATII RNA accumulation because depletion of HSATII RNA using ASOs in dox-pulsed iDUX4 cells completely abolished YTHDC1 nuclear aggregates (Fig. 4I); where only 2% of HSATII RNA-depleted cells contained YTHDC1 nuclear aggregates compared to 20% of control depleted cells (Fig. 4L). Furthermore, DUX4-expression or HSATII depletion did not impact m^6^A-related protein levels (Fig. S2), indicating that HSATII RNA affected protein localization and not protein stability.

Related with the sequestration of m^6^A-related factors, there was an increase in signal overlap between HSATII RNA aggregates and nuclear m^6^A signal (1873 ± 605) compared to ROI within the nucleoplasm (1583 ± 405) within dox-pulsed iDUX4 HSATII+ cells (Fig 4M and N). Interestingly, there was an overall increase in nuclear m^6^A signal within DUX4-expressing cells regardless of HSATII RNA accumulation (HSATII neg: 1551 ± 465, HSATII pos: 1801 ± 611) compared to uninduced control cells (950 ± 74) (Fig. 4O). Additionally, RNA dot blot probed using an m^6^A antibody showed a similar increase in total m^6^A RNA levels in dox-induced iDUX4 cells compared to uninduced controls (Fig. 4P). These data indicate that HSATII RNA associates with m^6^A-related factors, including m^6^A writer complex components (VIRMA and WTAP) and m^6^A-readers (YTHDC1), and their nuclear aggregation in DUX4-expressing cells is dependent on HSATII RNA accumulation. Furthermore, DUX4-expressing muscle cells have overall increased m^6^A RNA methylation levels suggesting dysregulation of RNA methylation processing.

### m^5^C-RNA methylation related factors are sequestered by HSATII RNA

To confirm localization of m^5^C RNA methylation factors with HSATII RNA, we performed immunofluorescence staining combined with HSATII RNA-FISH. RNA methyltransferase NSUN2 co-localized with HSATII RNA foci (Fig. 5A), where 67% of nuclei with NSUN2 nuclear aggregates in dox-pulsed iDUX4 cells showed co-localization with HSATII RNA foci (Fig. 5B). Nuclear NSUN2 signal was specifically enriched within HSATII RNA foci (2394 ± 165) compared to ROI within the nucleoplasm (2194 ± 93) (Fig. 5C). In connection, immunostaining using an anti-m^5^C antibody showed overlap of m^5^C signal with HSATII RNA foci (Fig. 5D), although multiple additional foci of m^5^C signal was also present. Overall, nuclear m^5^C signal was increased in dox-pulsed iDUX4 cells regardless of HSATII RNA accumulation (HSATII neg: 329 ± 112, HSATII pos: 313 ± 82) compared to uninduced cells (196 ± 40) (Fig. 5E; Fig. S3). However, within HSATII+ cell nuclei, m^5^C signal was enriched in HSATII RNA foci (447 ± 211) compared with ROI within the nucleoplasm (315 ± 135), and this enrichment was abolished when cells were treated with methylase inhibitor 5-azacytidine (5-azaC) (337 ± 172) (Fig. 5D and F).

**Figure 5.**
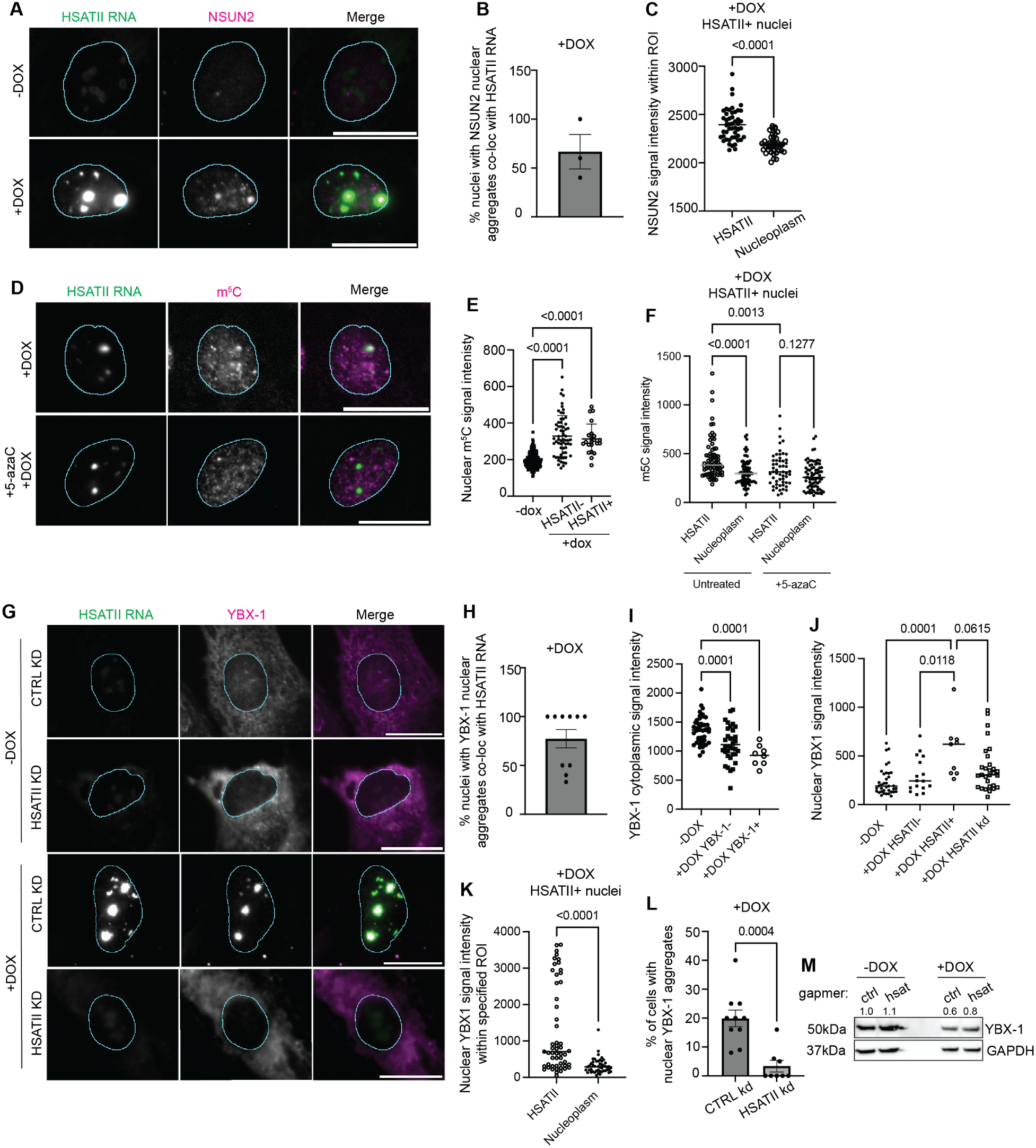
HSATII RNA sequesters m^5^C-related factors NSUN2 and YBX-1. (**A**) Combined immunofluorescence and HSATII RNA-FISH of HSATII (green) and NSUN2 (magenta) in –DOX or +DOX iDUX4 cells fixed at 24-hour time point. Scale bar = 20μm. Images are representative of three independent experimental replicates performed. (**B**) Fraction of nuclei with NSUN2 signal that at least have partial signal overlap with HSATII RNA foci in +DOX iDUX4 cells. N ≥ 200 nuclei. Dots represent mean of three independent experimental replicates. Data represent means ± SD. (**C**) Mean signal intensity of NSUN2 within specified ROI in the nucleus: either within HSATII RNA foci or the remainder of the nucleoplasm in +DOX HSATII+ nuclei in iDUX4 cells. Dots are each individual ROI. N ≥ 50 ROI. ROI within the nucleoplasm is drawn with relatively the same circumference as HSATII RNA foci ROI. Data are representative of three independent experimental replicates. Data represent means. Statistical differences between groups were analyzed employing Unpaired t-test. (**D**) Combined immunofluorescence and HSATII RNA-FISH of HSATII (green) and m^5^C (magenta) in +DOX iDUX4 cells either pre-treated with DMSO or 5µM 5-azaC for 24-hours, then dox-induced and fixed 24 hours post-induction. Scale bar = 20μm. Images are representative of three independent experimental replicates performed. (**E**) Mean signal intensity of nuclear m^5^C signal in –DOX or +DOX HSATII negative (HSATII-) or HSATII positive (HSATII+) cells fixed at 24-hour time point. Dots are individual nuclei. N ≥ 50 nuclei. Data are representative of three independent experimental replicates. Data represent means. Statistical differences between groups were analyzed employing One-way ANOVA Tukey’s multiple comparison test. (**F**) Mean signal intensity of m^5^C within specified ROI in the nucleus: either within HSATII RNA foci or the remainder of the nucleoplasm in untreated or 5-azaC treated +DOX iDUX4 cells. Dots are each individual ROI. N ≥ 55 ROI. ROI within the nucleoplasm is drawn with relatively the same circumference as HSATII RNA foci ROI. Data are representative of three independent experimental replicates. Data represent means. Statistical differences between groups were analyzed employing one-way ANOVA Tukey’s multiple comparison test. (**G**) Combined immunofluorescence and HSATII RNA-FISH of HSATII (green) and YBX-1 (magenta) in –DOX or +DOX iDUX4 cells with CTRL KD or HSATII KD and fixed 24-hours post-induction. Scale bar = 20μm. Images are representative of three independent experimental replicates performed. (**H**) Fraction of nuclei with YBX-1 signal that at least have partial signal overlap with HSATII RNA foci in +DOX iDUX4 cells. N ≥ 200 nuclei. Dots represent fields taken from representative experiment. Data represent means ± SD. (**I**) Cytoplasmic signal intensity of YBX-1 in –DOX or +DOX iDUX4 cells that have no YBX-1 nuclear aggregates (YBX-1-) or with YBX-1 nuclear aggregates (YBX-1+). N ≥ 200 nuclei. Dots are individual cells. Data represent means ± SD. Statistical differences between –DOX (control) and +DOX iDUX4 cells were analyzed employing one-way ANOVA Tukey’s multiple comparison test. (**J**) Nuclear mean signal intensity of YBX-1 within –DOX, +DOX HSATII-, +DOX HSATII+ or +DOX HSATII KD iDUX4 nuclei. Dots indicate individual nuclei. N = 10-40. Data are representative of three independent experimental replicates. Statistical differences between –DOX (control) and +DOX iDUX4 cells were analyzed employing one-way ANOVA Tukey’s multiple comparison test. (**K**) Mean signal intensity of YBX-1 within specified ROI in the nucleus: either within HSATII RNA foci or the remainder of the nucleoplasm in +DOX HSATII+ iDUX4 cells. Dots are each individual ROI. N ≥ 50 ROI. ROI within the nucleoplasm is drawn with relatively the same circumference as HSATII RNA foci ROI. Data are representative of three independent experimental replicates. Data represent means. Statistical differences between groups were analyzed employing Mann Whitney test. (**L**) Percent of cells with YBX-1 nuclear aggregates in +DOX iDUX4 cells with CTRL KD or HSATII KD. N ≥ 200 nuclei. Dots represent fields taken from representative experiment. Data represent means ± SD. Data are representative of three independent experimental replicates. Statistical differences between groups were analyzed employing Unpaired t-test. (**M**) Immunoblot of whole cell lysate from –DOX or +DOX iDUX4 cells with CTRL KD or HSATII KD and harvested at 24-hours post-induction. Blot was probed for YBX-1 total protein levels. GAPDH was used as loading control. Quantification of YBX-1 relative protein levels are indicated above YBX-1 blot.

The co-localization of NSUN2 and overlap of m^5^C signal with HSATII RNA suggested that HSATII may associate with m^5^C-reader proteins. Based on our proteomics, m^5^C-reader YBX-1 was identified as an HSATII-associated protein. Using immunofluorescence combined with HSATII RNA-FISH we confirmed that YBX-1 strongly co-localized with HSATII RNA foci in DUX4-expressing cells (Fig. 5G), where 77% of nuclei with nuclear YBX-1 aggregates in dox-pulsed iDUX4 cells showed co-localization with HSATII RNA foci (Fig. 5H). This was quite striking, as YBX-1 is a mainly cytoplasmic protein, yet YBX-1 cytoplasmic signal was significantly reduced in dox-pulsed iDUX4 cells that contained YBX-1 nuclear aggregates (926 ± 177) compared to uninduced control cells (1365 ± 233) (Fig. 5G and I). Nuclear YBX-1 signal was only present in dox-pulsed iDUX4 HSATII+ cells (563 ± 287) compared to uninduced (234 ± 138), dox-pulsed iDUX4 HSATII-cells (304 ± 182), or dox-pulsed HSATII KD iDUX4 cells (372 ± 235) (Fig. 5J). Within induced iDUX4 HSATII+ cells, YBX-1 signal was strongly enriched in HSATII RNA foci (1283 ± 1196) compared to ROI within the nucleoplasm (317 ± 208) (Fig. 5K). Nuclear aggregation of YBX-1 was dependent on HSATII RNA accumulation because depletion of HSATII RNA using ASOs in dox-pulsed iDUX4 cells completely abolished YBX-1 nuclear signal (Fig. 5G); where only 3% of HSATII RNA-depleted cells contained YBX-1 nuclear aggregates compared to 20% of control depleted cells (Fig. 5L). HSATII RNA-depletion did not affect total YBX-1 protein levels in dox-pulsed iDUX4 cells (Fig. 5M), indicating that HSATII RNA accumulation impacts YBX-1 localization and not protein stability. These data indicate that m^5^C-related factors, NSUN2 and YBX-1, co-localize and associate with intranuclear HSATII RNA. Additionally, YBX-1 nuclear aggregation is dependent on HSATII RNA accumulation.

### NSUN2 activity is necessary for YBX-1 localization with HSATII RNA

We hypothesized that NSUN2 activity may be necessary for YBX-1 association with HSATII RNA, as both were found to interact with and localize to HSATII RNA foci. NSUN2 has been shown to methylate noncoding RNA which impacts its processing, activity and turnover (Blanco et al., 2014; Hussain et al., 2013). First, we determined whether methylation contributed to the localization of YBX-1 with HSATII RNA. Dox-pulsed iDUX4 cells were treated with dimethyl sulfoxide (DMSO) as control or with 5-azaC for 20 hours and analyzed by microscopy for localization of YBX-1 and HSATII RNA (Fig. 6A). Treatment with 5-azaC significantly decreased the frequency of dox-pulsed iDUX4 HSATII+ cells that showed HSATII RNA foci overlap with nuclear YBX-1 aggregates from 24% in DMSO-treated to 5% in 5-azaC-treated cells (Fig. 6A and B). To determine whether enrichment of m^5^C signal with HSATII RNA was NSUN2 dependent, we depleted uninduced or dox-pulsed iDUX4 cells of NSUN2 using small interfering RNAs (siRNA) or non-targeting control siRNA and performed immunofluorescence combined with HSATII RNA-FISH to determine signal co-localization (Figs. 6C and D; and S4A and B). Depletion of NSUN2 did not drastically alter total nuclear m^5^C signal (Fig. S4C and D), however in dox-pulsed iDUX4 HSATII+ cells, NSUN2 depletion significantly decreased m^5^C signal intensity within HSATII RNA foci (183 ± 98) compared to control siRNA-depleted cells (281 ± 144) (Fig. 6C and D), suggesting that NSUN2 may impact HSATII RNA methylation.

**Figure 6.**
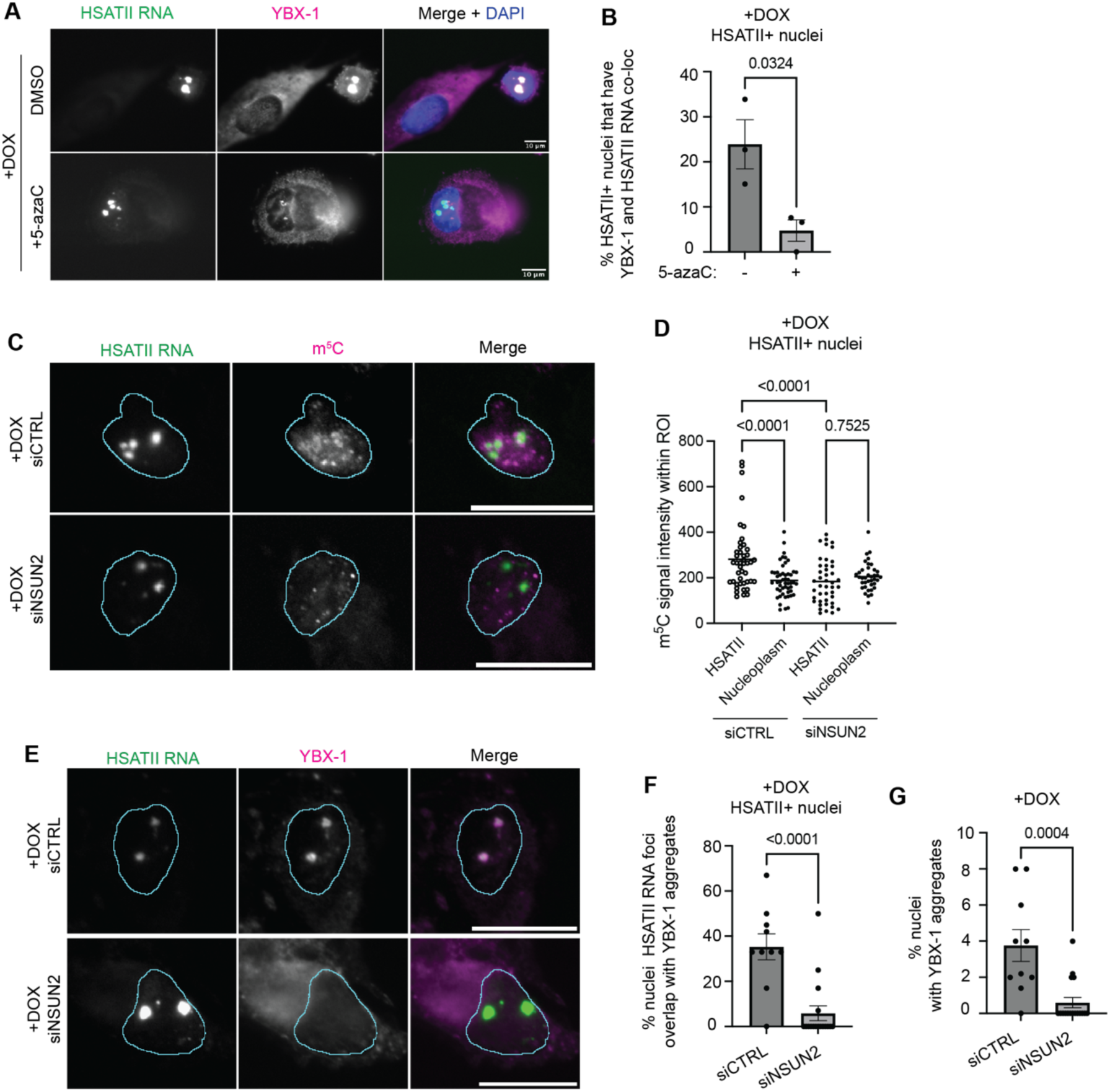
NSUN2 activity is necessary for YBX-1 association with HSATII RNA. (**A**) Combined immunofluorescence and HSATII RNA-FISH of HSATII (green) and YBX-1 (magenta) in +DOX iDUX4 cells either pre-treated with DMSO or 5µM 5-azaC for 24-hours, then dox-induced and fixed 24 hours post-induction. Scale bar = 10μm. Images are representative of three independent experimental replicates performed. (**B**) Percent of HSATII+ nuclei with YBX-1 signal that overlap with HSATII RNA foci in +DOX iDUX4 cells with or without 5-azaC treatment. N ≥ 200 nuclei. Dots represent mean of independent experimental replicates. Data represent means ± SD. Statistical differences between groups were analyzed employing two-tailed Unpaired t-test. (**C**) Combined immunofluorescence and HSATII RNA-FISH of HSATII (green) and m^5^C (magenta) in +DOX iDUX4 cells either pre-treated with siRNAs targeting control sequences (siCTRL) or NSUN2 (siNSUN2) for 24-hours, then dox-induced and fixed 24 hours post-induction. Scale bar = 20μm. Images are representative of three independent experimental replicates performed. (**D**) Mean signal intensity of m^5^C within specified ROI in the nucleus: either within HSATII RNA foci or the remainder of the nucleoplasm in +DOX HSATII+ iDUX4 cells with control (siCTRL) or NSUN2 (siNSUN2) depletion. Dots are each individual ROI. N ≥ 50 ROI. ROI within the nucleoplasm is drawn with relatively the same circumference as HSATII RNA foci ROI. Data are representative of three independent experimental replicates. Data represent means. Statistical differences between groups were analyzed employing one-way ANOVA Tukey’s multiple comparison test. (**E**) Combined immunofluorescence and HSATII RNA-FISH of HSATII (green) and YBX-1 (magenta) in +DOX iDUX4 cells either pre-treated with siRNAs targeting control sequences (siCTRL) or NSUN2 (siNSUN2) for 24-hours, then dox-induced and fixed 24 hours post-induction. Scale bar = 20μm. Images are representative of three independent experimental replicates performed. (**F**) Percent of HSATII+ nuclei with YBX-1 signal that overlap with HSATII RNA foci in +DOX iDUX4 cells with control (siCTRL) or NSUN2 (siNSUN2) depletion. N ≥ 200 nuclei. Dots represent fields taken from representative experiment. Data represent means ± SD. Statistical differences between groups were analyzed employing Mann Whitney test. (**G**) Percent of nuclei with YBX-1 nuclear aggregates in +DOX iDUX4 cells with control (siCTRL) or NSUN2 (siNSUN2) depletion. N ≥ 200 nuclei. Dots represent fields taken from representative experiment. Data represent means ± SD. Data are representative of three independent experimental replicates. Statistical differences between groups were analyzed employing Mann Whitney test.

Next, we determined whether NSUN2 activity was necessary for YBX-1 co-localization with HSATII RNA foci. NSUN2-specific methylation of RNA has been shown to recruit YBX-1 for RNA regulation (Chen et al., 2024; Liu et al., 2024; Zhang et al., 2024). Nuclear localization of YBX-1 was completely diminished in NSUN2-depleted dox-pulsed iDUX4 HSATII+ cells compared to control siRNA-treated cells (Fig. 6E). In dox-pulsed iDUX4 HSATII+ cells treated with control siRNAs, 35% of nuclei showed HSATII RNA foci and YBX-1 nuclear signal overlap, whereas this co-localization decreased to only 6% in NSUN2-depleted cells (Fig. 6F). Furthermore, there was an overall decrease in frequency of cells with YBX-1 nuclear aggregates, from 4% in control siRNA treated dox-pulsed iDUX4 cells to less than 1% in NSUN2-depleted cells (Fig. 6G). NSUN2 depletion did not impact total YBX-1 protein levels (Fig. S4A and B), suggesting that NSUN2 activity influenced YBX-1 localization and not protein homeostasis. These data point to NSUN2 activity being necessary for YBX-1 recruitment to HSATII RNA foci in DUX4-expressing cells.

### YBX-1 specifically localizes with HSATII double-stranded RNA

Our published work shows that DUX4 expression in skeletal muscle induces temporally-regulated, bidirectional transcription of HSATII sequences which leads to HSATII single-stranded RNA (“ssRNA”; 12 hour post-dox) and double-stranded RNA (“dsRNA”; 20 hour post-dox) accumulation (Shadle et al., 2019). Single-stranded and double-stranded HSATII RNA have been shown to cause protein aggregation of RBPs including eIF4A3 and ADAR1 (Shadle et al., 2019). Bidirectional transcription and temporal expression of pericentric satellites is not unique to DUX4-expressing myoblast cells, but also observed in mouse (Probst et al., 2010) and human (Yandim and Karakulah, 2019) development, where mouse Dux and human DUX4 are briefly expressed. In human myoblast cells, DUX4 expression leads to the production of HSATII “reverse” transcripts at 12 hours post-dox induction which form nuclear ssRNA aggregates, followed by production of HSATII “forward” transcripts 16 hours post-dox induction, and the formation of dsRNA foci (Shadle et al., 2019). This suggests potential differential regulatory roles of HSATII depending on HSATII RNA structure, i.e., single or double stranded, and we considered whether this played a role in regulating protein aggregation in DUX4-expressing myoblast cells.

Even though nearly 80% of nuclei that contained YBX-1 nuclear aggregates in dox-pulsed iDUX4 cells showed localization with HSATII RNA foci (see Fig. 5H), only 16% of HSATII+ nuclei showed HSATII and YBX-1 co-localization (Fig. 7A and C), indicating that not all HSATII RNA associated with or sequestered YBX-1. We postulated that HSATII RNA structure dictated selective interaction with YBX-1. DUX4 robustly induces accumulation of intranuclear dsRNA, and these dsRNA sequences were enriched for HSATII repeats (Shadle et al., 2019). We have previously shown that the dsRNA antibody “K1” shows strict localization with HSATII dsRNA foci and depletion of HSATII RNA abolishes K1 signal in DUX4-expressing cells (Shadle et al., 2019), indicating that K1 signal is a good readout for HSATII dsRNA. To resolve the association of YBX-1 with HSATII ssRNA or dsRNA we performed a time course experiment where we dox-pulsed iDUX4 cells and then fixed cells at either 16 hours (12 hours post dox washout) to capture HSATII ssRNA detected by the FISH probe to the initially expressed “reverse” strand, or 24 hours (20 hours post-dox induction) to capture the HSATII dsRNA using the K1 antibody. At the 16-hour time point only the probe targeting HSATII reverse transcripts showed signal for HSATII RNA (8% cells HSATII ssRNA positive) and the K1 antibody did not show any signal, indicating no accumulation of HSATII dsRNA (Fig. 7A and B). By the 24-hour time point 21% of cells had HSATII foci detected by the reverse strand probe (ssRNA and/or dsRNA) and 6% had K1 foci (dsRNA) (Fig. 7A and B). HSATII+ nuclei showed no nuclear aggregation or localization of YBX-1 with HSATII RNA foci at the 16-hour time point. At the 24-hour time point roughly 16% of HSATII+ nuclei had YBX-1 nuclear aggregation and co-localization (Fig. 7A and C) and 70% of dsRNA+ nuclei had nuclear aggregation and co-localization of YBX-1 with dsRNA foci (Fig. 7A and D). Nuclear YBX-1 signal was strongly localized within dsRNA foci (1249 ± 973) compared to ROI within the nucleoplasm (402 ± 557) in dox-pulsed iDUX4 dsRNA+ nuclei (Fig. 7E). These data indicate that YBX-1 is primarily sequestered by HSATII dsRNA.

**Figure 7.**
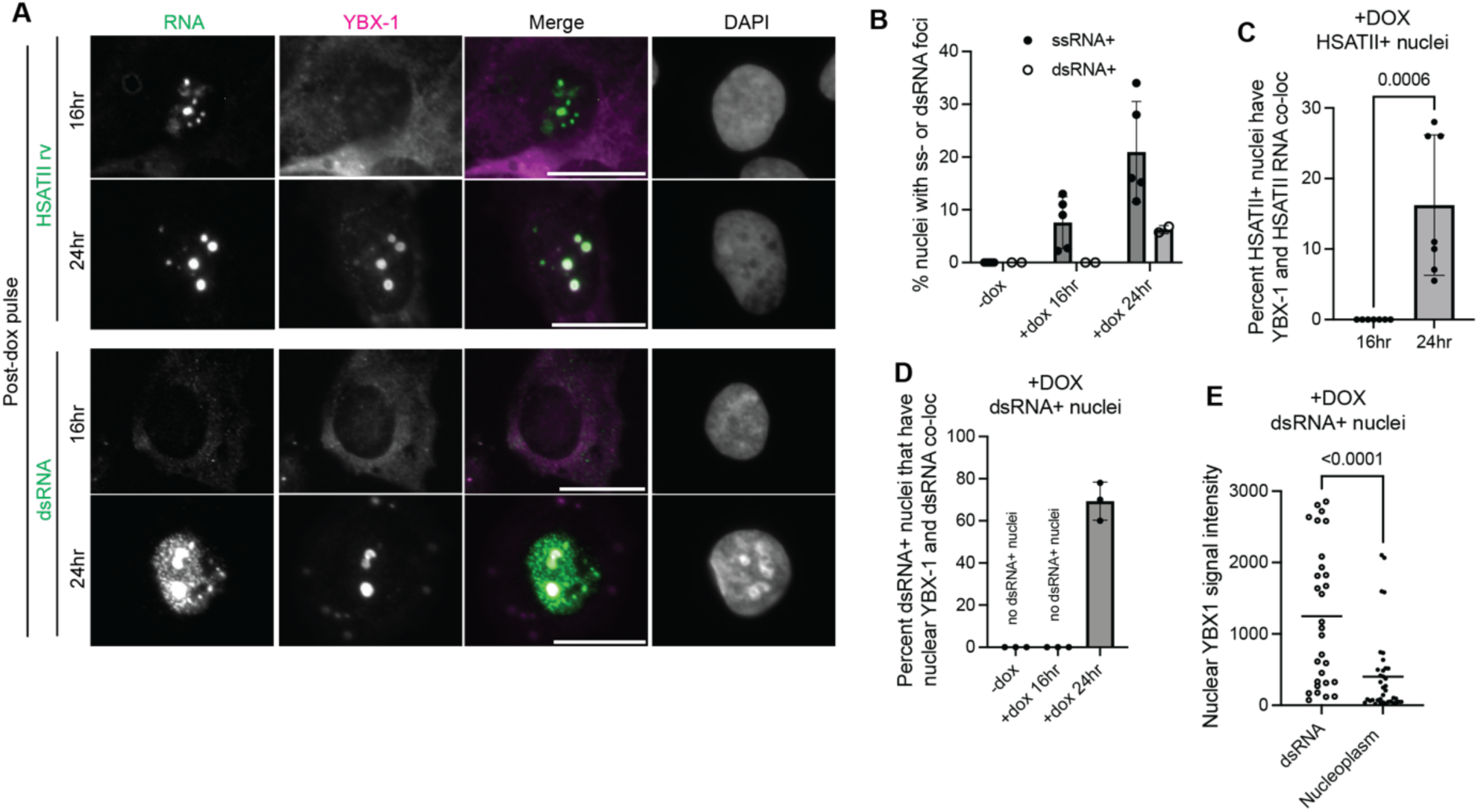
YBX-1 associates with HSATII dsRNA. (**A**) Top panel: combined immunofluorescence and HSATII RNA-FISH of HSATII (green) and YBX-1 (magenta) in +DOX iDUX4 cells and fixed at either 16-hour time point or 24-hour time point; Bottom panel: Immunofluorescence of dsRNA (green) and YBX-1 (magenta) in +DOX iDUX4 cells and fixed at either 16-hour time point or 24-hour time point. Scale bar = 20μm. Images are representative of three independent experimental replicates performed. (**B**) Frequency of nuclei with either HSATII ssRNA or dsRNA in –DOX or +DOX iDUX4 cells at either 16-hour time point or 24-hour time point. N ≥ 200 nuclei. Dots represent independent experimental replicates. Data represent means ± SD. (**C**) Percent of HSATII+ nuclei with YBX-1 signal that overlap with HSATII RNA foci in +DOX iDUX4 cells at either 16-hour time point or 24-hour time point. N ≥ 200 nuclei. Dots represent fields taken from representative experiment. Data represent means ± SD. Statistical differences between groups were analyzed employing Mann Whitney test. (**D**) Percent of dsRNA+ nuclei with YBX-1 signal that overlap with dsRNA foci in +DOX iDUX4 cells at either 16-hour time point or 24-hour time point. N ≥ 200 nuclei. Dots represent independent experimental replicates. Data represent means ± SD. (**E**) Mean signal intensity of YBX-1 within specified ROI in the nucleus: either within dsRNA foci or the remainder of the nucleoplasm in +DOX dsRNA+ iDUX4 cells. Dots are each individual ROI. N ≥ 30 ROI. ROI within the nucleoplasm is drawn with relatively the same circumference as dsRNA foci ROI. Data are representative of three independent experimental replicates. Data represent means. Statistical differences between groups were analyzed employing two-tailed Unpaired t-test.

### HSATII-RNP complexes impact mRNA processing pathways

We have previously shown that HSATII expression impacts localization and function of nuclear regulatory proteins which has severe consequences on cellular pathway regulation (Arends et al., 2024). HSATII RNA accumulation in DUX4-expressing cells impacts the localization of several canonical RNA processing factors (see Figs. 4 and 5) (Shadle et al., 2019). Our work, as well as others, has demonstrated that DUX4 expression in muscle cells impacts RNA processing pathways including nonsense mediated decay (NMD), splicing and RNA methylation (see Fig. 4M) (Campbell et al., 2023; Geng et al., 2012; Jagannathan et al., 2019; Snider et al., 2010; Young et al., 2013). We hypothesized that HSATII sequestration of RNA binding proteins may influence RNA processing pathways. We have previously demonstrated that HSATII RNA sequesters exon junction complex (EJC) component eIF4A3 (Shadle et al., 2019), and the EJC, as well as eIF4A3, have been implicated in affecting splicing and NMD (Le Hir et al., 2016; Ryu et al., 2019). Furthermore, RNA methylation and related factor activity impact alternative RNA processing events and NMD (Elvira-Blazquez et al., 2024; Li et al., 2019; Liu et al., 2021; Xiao et al., 2016; Yang et al., 2019; Yue et al., 2018).

To interrogate whether HSATII-RNP formation impacts RNA processing events downstream of DUX4-expression, we performed poly(A)-selected, short-read RNA-sequencing on uninduced and dox-pulsed iDUX4 cells treated with HSATII-specific ASOs or control ASOs (Fig. S5A). Depletion of HSATII RNA from DUX4-expressing cells did not significantly impact DUX4-target gene expression (Suppl. Table 3). However, depletion of HSATII RNA did influence the alternative splicing events mediated by DUX4 expression. Over 500 genes had statistically significant differential splicing events between control and HSATII depleted samples in DUX4-expressing cells (Suppl. Table 3). Significant differential splicing events resulted in 5’ UTR gain/loss, intron retention, and alternative transcriptional termination site usage within DUX4-expressing cells with HSATII depletion (Fig. 8A). Gene set enrichment using Enrichr (Chen et al., 2013; Kuleshov et al., 2016; Xie et al., 2021) found that the these genes were enriched in biological processes including regulation of innate immune response, telomere regulation, protein transport, apoptotic signaling in response to oxidative stress, RNA methylation, and double-stranded break repair (Fig. 8B).

**Figure 8.**
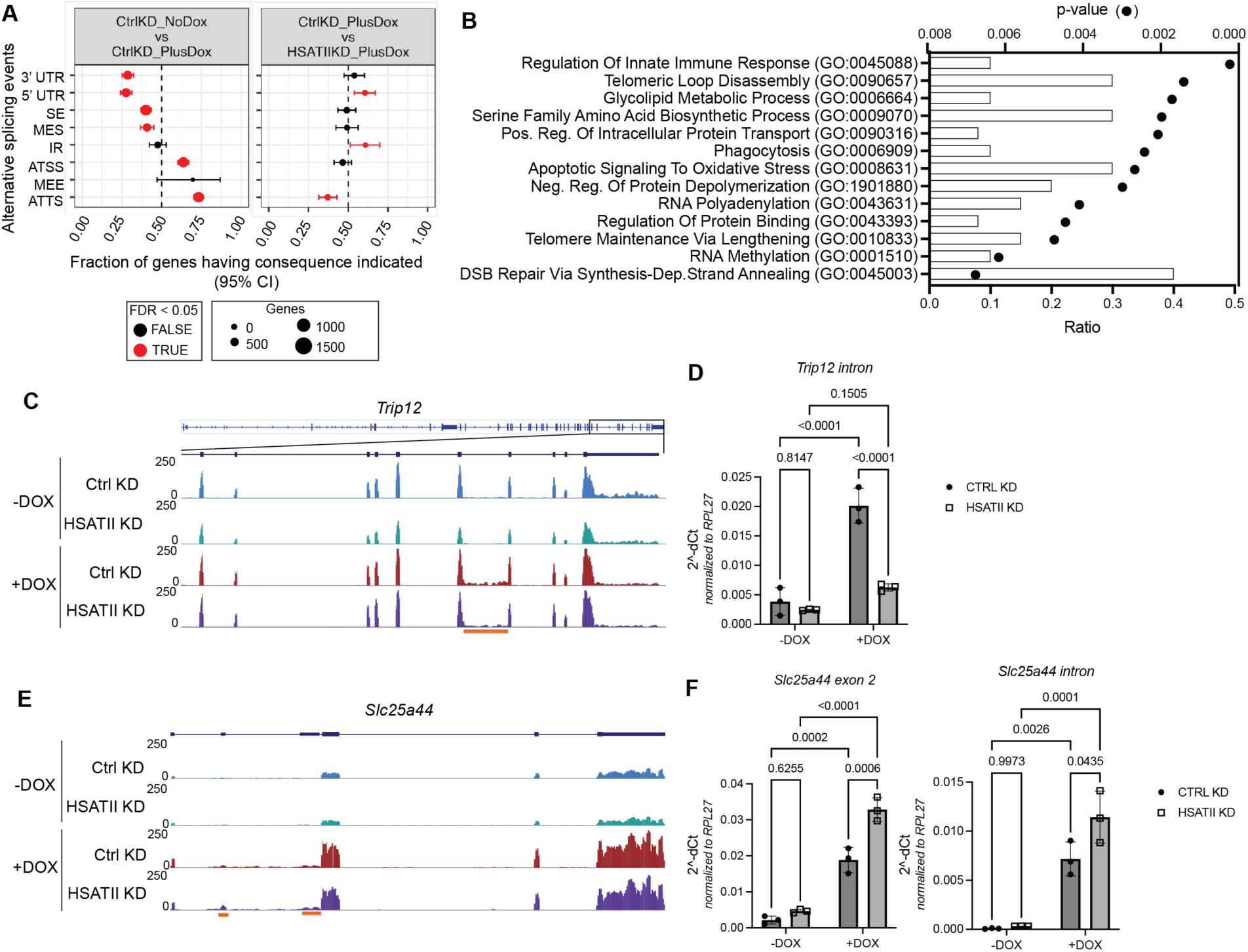
HSATII-RNP complexes impact RNA splicing in DUX4-expressing cells. (**A**) RNA processing consequences and fraction of genes with indicated consequence in control depleted (CTRL) –DOX or +DOX iDUX4 cells or in +DOX iDUX4 cells with control (CTRL KD) or HSATII depletion (HSATII KD). RNA-sequencing was performed in biological duplicate. UTR: untranslated region; SE: skipped exon; IR: intron retention; MES: mutually exclusive splicing; ATSS: alternative transcription start site; MEE: mutually exclusive exon; ATTS: alternative transcription termination site; FDR: false discovery rate. (**B**) Top enrichment of Biological Processes of significantly affected genes between +DOX iDUX4 CTRL KD compared to HSATII KD. Ratio indicated number of genes within pathway. (**C**) UCSC genome browser tracks showing RNA-seq reads which map to gene *Trip12* from –DOX iDUX4 cells with Ctrl (blue) or HSATII-depletion (green) or +DOX iDUX4 cells with Ctrl (red) or HSATII-depletion (purple). +DOX Ctrl cells have increased intron retention (highlighted region, orange bar). HSATII depletion decreases intron retention. (**D**) Expression of *Trip12* with intron retention in –DOX or +DOX iDUX4 cells with (HSATII KD) or without (CTRL KD) HSATII depletion. N= 3. Data represent means ± SD. Statistical differences between groups were analyzed employing 2-way ANOVA Tukey’s multiple comparison test. (**E**) UCSC genome browser tracks showing RNA-seq reads which map to gene *Slc25a44* from –DOX iDUX4 cells with Ctrl (blue) or HSATII-depletion (green) or +DOX iDUX4 cells with Ctrl (red) or HSATII-depletion (purple). +DOX Ctrl cells have decreased exon 2 and intron inclusion (highlighted region, orange bar). HSATII depletion increases exon 2 retention and intron inclusion. (**F**) Expression of *Slc25a44* containing exon 2 (left panel) or intron retaining (right panel) isoforms –DOX or +DOX iDUX4 cells with (HSATII KD) or without (CTRL KD) HSATII depletion. N= 3. Data represent means ± SD. Statistical differences between groups were analyzed employing 2-way ANOVA Tukey’s multiple comparison test.

Of particular interest, we validated differential RNA isoform usage of *Trip12*, an E3 ubiquitin ligase that modulates DNA damage meditated ubiquitin signaling (Gudjonsson et al., 2012) and cell cycle (Chen et al., 2010; Kajiro et al., 2011). We have previously shown that DUX4 expression impacts cell cycle progression and HSATII transcription impedes PRC1-mediated ubiquitination at sites of DNA damage (Arends et al., 2024). Therefore, we interrogated whether HSATII expression could impact DNA damage response factors through RNA splicing regulation. HSATII-RNPs promoted intron retention within *Trip12* in DUX4-expressing cells (Fig. 8C and D), where upon depletion of HSATII RNA there was a significant loss of *Trip12* intron retention. Additionally, HSATII-RNPs impacted RNA splicing of *Slc25a44*, a mitochondrial branched-chain amino acid carrier (Yoneshiro et al., 2019). DUX4-expression in muscle cells has been shown to impact mitochondrial function (Heher et al., 2022; Zhang et al., 2022). HSATII-RNPs promoted both exon and intron exclusion of *Slc25a44*, where depletion of HSATII RNA resulted in increased exon retention and intron inclusion in DUX4-expressing cells (Fig. 8E and F). Furthermore, HSATII-RNPs promoted differential isoform expression of *eIF4ENIF1*, a translational regulator that interacts with eIF4E (Ferraiuolo et al., 2005). We have previously reported that DUX4 expression results in broad translational repression in muscle cells, partially through the regulation of eIF4E activity (Hamm et al., 2023). HSATII-RNPs impacted the differential isoform usage of *eIF4ENIF1* (specifically canonical isoform ENST00000330125.10 and alternative isoform ENST00000423097.5) when analyzed using pairwise comparisons using Isoform Switch AnalyzeR (Vitting-Seerup and Sandelin, 2019) (Fig. S5B) and validated by RT-qPCR using isoform-specific primers (Fig. S5C). Immunoblot of eIF4ENIF1 also detected differential protein isoforms that correlated with the predicted translated protein product of the specific *eIF4ENIF1* RNA isoform (ENST00000330125.10, predicted protein product ∼100kDa; ENST00000423097.5, predicted protein product ∼20kDa) (Fig. S5D). Our data indicate that HSATII-RNP formation impacts RNA processing pathways, leading to differential splicing of genes downstream of DUX4 expression which are implicated in FSHD pathology.

## DISCUSSION

In this study, we uncovered that DUX4 expression leads to the stable, long-lived accumulation of nucleolar-associated RNA and HSATII RNA. These stable RNA aggregates sequester RBPs and cause nuclear protein aggregation of proteins known to be impacted by DUX4 expression, TDP-43 and SC35, as well as newly identified proteins, m^6^A and m^5^C RNA methylation related factors. Formation of HSATII-YBX-1 RNP complexes is dependent on HSATII double-stranded RNA (dsRNA) accumulation and NSUN2 methylation activity. Importantly, aberrant formation of HSATII-RNP complexes in muscle cells affects RNA processing downstream of DUX4 expression. Our study implicates HSATII RNA and the formation of HSATII-RNP complexes as mediators of gene regulation.

An important finding in this study is the induction of stable RNA aggregates by DUX4. We found that major components of these stable RNA aggregates were nucleolar-associated RNAs and HSATII RNA. Our previous work indicated that DUX4 induces accumulation of dsRNA and particularly HSATII RNA, where DUX4 binds to and transcriptionally activates HSATII (Shadle et al., 2019; Young et al., 2013). However, the mechanism driving the apparent accumulation of nucleolar-associated RNA, including ribosomal RNA (rRNA) is less clear. Loss of rDNA stability has been shown to induce age-dependent aggregation of rRNA-binding proteins through aberrant overproduction of rRNAs (Paxman et al., 2022). DUX4 expression induces constitutive DNA damage and oxidative stress which could impact rDNA integrity leading to aberrant overexpression of rRNA genes (Arends et al., 2024; Dmitriev et al., 2016).

The apparent reorganization of the nucleolus and mis-localization of nucleolar proteins, including FBL and NPM1, suggest that DUX4 may impact nucleolar compartmentalization. Our data indicate that nucleolar disruption co-occurs in nuclei that contain stable intranuclear RNA aggregates (see Fig. 1L). During the stages of early embryogenesis when Dux/DUX4 is expressed, nucleolus-precursor bodies (NPB) exist, but not mature nucleolus; however, at the end of the 2-cell stage in mouse or 4-cell stage in human after Dux/ DUX4 expression occurs, respectively, rRNA gene synthesis occurs, followed by maturation of the NPB into the nucleolus (Kresoja-Rakic and Santoro, 2019). Interestingly, pericentromeric repeats surround the NPB and their spatial positioning around the NPB is necessary for their heterochromatin silencing within the embryo (Jachowicz et al., 2013). Upon transcriptional activation of pericentromeric satellite repeats during the 2-cell/ 4-cell stage, heterochromatin formation occurs, and satellites associate with NPBs and structural changes occur. Our data suggest that DUX4-induction of HSATII expression in muscle cells could indirectly impact nucleolar structure by disrupting RNA metabolism. Interestingly, 2-cell (2C) mouse embryos and 2C-like cells that emerge from mouse embryonic stem cell cultures, exhibit circular immature nucleoli which is similar to what we observe in DUX4-expressing myoblast cells (Xie et al., 2022) (Fig. 1J and K). However, the direct impact of HSATII expression on nucleolar architecture in DUX4 expressing cells still needs to be determined.

This study highlights a mechanism driving protein aggregation in DUX4-expressing muscle cells, via protein sequestration through induced stable intranuclear RNA accumulation. Specific RNAs promote the formation of RNP complexes by acting as scaffolds for RBPs. Evidence indicates that highly contacted RNAs are structured, have long UTRs, or are highly repetitive (Cid-Samper et al., 2018). HSATII is a tandem repetitive element that contains several repeat units within an RNA molecule. This highly repetitive RNA may in turn cause specific protein sequestration. Indeed, we found that HSATII RNA sequesters specific RBPs including RNA methylation related factors and TDP-43, among others. Aggregation of other proteins observed in DUX4-expressing muscle cells may be due to accumulation of other RNAs including nucleolar RNAs, as shown with EU+ nucleolar RNA and SC35 in this study. However, other mechanisms may be driving protein aggregation in DUX4-expressing muscle cells including dysregulation of RNA surveillance pathways. Defective mRNA surveillance pathways have been reported to drive protein production leading to protein aggregation (Jamar et al., 2018). Recent work has shown that dysregulation of RNA quality control pathways in the context of DUX4 expression results in aberrant protein production and protein aggregates (Campbell et al., 2023). However, this mechanism does not account for all protein aggregation in FSHD muscle cells and thus our current study underscores another mechanism driving aberrant protein aggregation in FSHD muscle cells.

Our work is the first to comprehensively identify endogenous HSATII RNA interacting proteins. The association with m^5^C and m^6^A RNA methylation factors was quite interesting, as satellite repeats have been associated with these modifications in development and disease (Duda et al., 2021; Ninomiya et al., 2021; Timcheva et al., 2022). RNA modifications have emerged as important posttranscriptional regulators of gene expression and RNA processing. Of such modifications, N^6^-methyladenosine (m^6^A) and 5-methylcytosine (m^5^C) have been shown to affect normal development by impacting splicing, export, stability and/or translation of RNA (Frye and Blanco, 2016; Roundtree et al., 2017; Shi et al., 2019). The biological functions of m^6^A and m^5^C are mediated by the writer and eraser which install and remove the methylation, respectively, and reader proteins which recognize it. Perturbations in these factors and/or aberrant expression has been linked to disease and cancer, suggesting a critical role for RNA modification in cellular function (Blanco et al., 2014; Roundtree et al., 2017; Shi et al., 2019; Sun et al., 2023). Thus, the aberrant association of RNA methylation factors with HSATII could inhibit their normal activity and lead to dysregulation in global RNA methylation. The mechanism of how HSATII sequestration of RNA methylation factors impact their activity still needs to be explored further.

Our work uncovered that HSATII RNA, specifically HSATII dsRNA, preferentially sequesters m^5^C reader protein YBX-1 through RNA accumulation and NSUN2 activity. Importantly, our study is the first to propose the dsRNA association of YBX-1. Why YBX-1 preferentially associates with HSATII is unclear. Future work will dissect the mechanisms regulating recruitment of factors to HSATII and how structure facilitates these interactions. Importantly, YBX-1 plays a critical part in myogenesis, and YBX-1 depletion was shown to impact differentiation in mouse myoblasts (Kobayashi et al., 2015). Sequestration of YBX-1 by HSATII dsRNA may in turn impact its localization with target transcripts necessary for promoting myoblast differentiation.

A proportion of the RNA processing defects identified in DUX4-expressing cells is a direct result of the formation of HSATII-RNP complexes. How HSATII sequestration of each factor impacts RNA processing is compounded by the fact that HSATII RNA sequesters several RBPs known to have roles in RNA processing. HSATII sequestration of m^6^A– and m^5^C-related factors may have profound effects on cellular RNA methylation. Indeed, we observed an increase in total m^6^A RNA methylation in DUX4-expressing cells. RNA methylation has known roles in regulating RNA processing including mRNA splicing, RNA localization and stability (Covelo-Molares et al., 2018; Sun et al., 2023). HSATII sequesters regulators of splicing and NMD including eIF4A3 which could contribute to the NMD pathway dysregulation observed in DUX4-expressing muscle cells. Future work will determine how HSATII sequestration of RNA methylation factors and other RBPs impact the RNA processing of specific genes. Importantly, genes impacted by HSATII-RNP formation are enriched in pathways that are dysregulated by DUX4 expression, including innate immune response signaling, telomere maintenance, cellular stress response, DNA damage and cell cycle regulation (Arends et al., 2024; Dmitriev et al., 2016; Hamm et al., 2023; Heher et al., 2022; Himeda and Jones, 2019; Spens et al., 2023; Xu et al., 2014). Through validation, we identified differential RNA splicing of *Trip12*, an E3 ubiquitin ligase, *Slc25a44*, a mitochondrial branched chain amino acid carrier, and *eIF4ENIF1*, a translation initiation factor, being mediated by HSATII-RNP. Differential RNA isoforms of these genes may lead to functional variability, potentially modifying their ability to interact with co-factors or impacting their function. Thus, HSATII-RNP formation impacts RNA processing pathways which may contribute to the dysregulation of pathways downstream of DUX4 expression in FSHD muscle cells.

## ACKNOWLEDGEMENTS

This research was supported by NIH NIAMS R01AR045203 (SJT); NCI T32CA009657 (TA); Friends of FSH Research (TA, SJT). We would also like to thank the Fred Hutchinson Cancer Center Cellular Imaging Shared Resource for assistance with the microscopy and image analysis, specifically Lena Schroeder and support through the Cellular Imaging Shared Resource (RRID:SCR_022609) of the Fred Hutch/ University of Washington/ Seattle Children’s Cancer Consortium (P30 CA015704).

## AUTHOR CONTRIBUTIONS

Author contributions: Conceptualization: T. Arends and S.J. Tapscott. Formal analysis: T. Arends and S.R. Bennett. Funding acquisition: T. Arends and S.J. Tapscott. Investigation: T. Arends. Project administration: T. Arends and S.J. Tapscott. Supervision: S.J. Tapscott. Validation: T. Arends. Visualization: T. Arends and S.R. Bennett. Writing – original draft: T. Arends. Writing – review & editing: T. Arends, S.R. Bennett and S.J. Tapscott.

## MATRIALS AND METHODS

### Cell culture

Immortalized MB135 myoblast cells (*H. sapiens*, female, Fields Center for FSHD and Neuromuscular Research at the University of Rochester Medical Center, https://www.urmc.rochester.edu/neurology/fshd-center.aspx) that contain a tet-inducible DUX4 transgene (iDUX4) (RRID:Addgene_99281) (Resnick et al., 2019) were cultured in F10 medium (Gibco/ ThermoFisher Scientific) supplemented with 20% Fetal Bovine Serum (GE Healthcare Life Sciences) and 1% penicillin/streptomycin (Gibco/ ThermoFisher Scientific), 10ng/mL recombinant human FGF (PeproTech) and 1μM dexamethasone (Sigma-Aldrich). iDUX4 cells were induced in the presence of 2μg/mL of doxycycline-hyclate (“DOX”; Sigma-Aldrich) for 4-hrs and assayed 20-hrs post-induction, unless otherwise noted.

### EU Click-iT immunofluorescence

Cells were cultured in 35mm dishes or grown in 4-or 8-well chamber slides and were assayed at various time points post-induction (24-, 48-, 72-, or 96-hrs post-induction). 0.1mM 5-ethynyl uridine (ThermoFisher Scientific) was incubated with cells after dox-induction for 16-hrs followed by a washout and then fixed at various time points. Cells were fixed in 4% paraformaldehyde (Electron Microscopy Sciences) for 10 min at RT, then quenched in 0.125M glycine for 5 min at room temperature (RT). Cells were permeabilized (0.5% Triton X-100 in 1X PBS) for 15 minutes at RT. EU-labeling click reaction solution was made using 0.25M conjugated-azide (biotin: Vector labs; AF488-conjugate: Sigma), 0.5M CuSO4, 0.25M THPTA, 0.1M sodium L-ascorbate in 1X PBS and incubated for 15 minutes at RT. Cells were washed with 1X PBS and incubated with 1 mL click reaction solution for 30 minutes at RT. Reaction was quenched 3x in 0.5% Triton X-100 in 1X PBS with 2mM EDTA for 3 minutes at RT. For combined antibody staining: Cells were then incubated in permeabilization and block buffer (0.5% Triton X-100, 5% Normal donkey serum (Jackson Immuno Research) for 1 hour at RT and subsequently incubated overnight at 4°C with primary antibody in staining buffer (0.1% Triton X-100, 1% BSA). Cells were washed with 1X PBS 15 min (2x), then incubated with secondary antibody for 1 hour at RT in staining buffer. Cells in 35mm dishes were washed with 1X PBS and nuclei were stained using DAPI (1:1000, Sigma-Aldrich) for 10 min at RT. Cells in chamber slides were briefly air-dried and mounted using ProLong Glass Antifade Mounting with NucBlue (ThermoFisher Scientific). Cells were imaged in 1X PBS on a Zeiss AxioPhot fluorescence microscope using 25X/0.80NA water immersion objective or oil at RT. All fluorescence channels were imaged at non-saturating levels, and settings were kept identical between all samples within replicates used for comparisons or quantifications. For antibodies used refer to “Antibodies, primers and reagents” section.

### Combined HSATII RNA-FISH and EU Click-iT immunofluorescence

Cells were cultured in 35mm dishes or grown in 4-or 8-well chamber slides. EU-click it immunofluorescence was performed as stated in “EU Click-iT immunofluorescence” section with modifications. After secondary staining cells were re-fixed in 4% paraformaldehyde for 7 min at RT. Cells were then washed in hybridization wash buffer (2X SSC, 50% Formamide) for 10 min at RT. Locked nucleic acid, FITC-conjugated HSATII probes were purchased from QIAGEN and are based on the sequence used in previous publications (Hall et al., 2017). Probe: 5′ FAM-ATTCCATTCAGATTCCATTCGATC detects the reverse HSATII transcript. HSATII probes were diluted to 5.0 pmol/mL in whole chromosome painting buffer (50% formamide (Sigma-Aldrich), 2X SSC (Invitrogen), 10% dextran sulfate and hybridized overnight at 37°C. Cells were washed with 15% formamide/2X SSC for 20 min at 37°C, 2X SSC for 20 min at 37°C and 2X SSC for 5 min at RT. Cells were washed with 1X PBS and nuclei were stained using DAPI for 10 min at RT. Cells in chamber slides were briefly air-dried and mounted using ProLong Glass Antifade Mounting with NucBlue (ThermoFisher Scientific). Cells in 35mm dishes were imaged in 1X PBS on a Zeiss AxioPhot fluorescence microscope using either 25X/0.80NA or 40X/0.90NA water immersion objective at RT. All fluorescence channels were imaged at non-saturating levels, and settings were kept identical between all samples within replicates used for comparisons or quantifications. For antibodies used refer to “Antibodies and primers” section.

### Immunofluorescence

Cells cultured in 35mm dishes were washed with 1X PBS then treated with cold Cytoskeletal (CSK) buffer (100mM NaCl, 300mM Sucrose, 3mM MgCl_2_, 10mM Pipes pH 6.8, 0.2mM Triton X-100) for 5 min. Cells were fixed in 2% or 4% paraformaldehyde (Electron Microscopy Sciences) for 10 min at RT, then quenched in 0.125M glycine for 5 min at RT. Cells were permeabilized and blocked (0.5% Triton X-100, 5% Normal donkey serum (Jackson Immuno Research) for 1 hour at RT and subsequently incubated overnight at 4°C with primary antibody in staining buffer (0.1% Triton X-100, 1% BSA). Cells were washed with 1X PBS 15 min (2x), then incubated with secondary antibody for 1 hour at RT in staining buffer. Cells were washed with 1X PBS and nuclei were stained using DAPI (1:1000, Sigma-Aldrich) for 10 min at RT. For m^6^A immunostaining: Cells were fixed in 4% paraformaldehyde overnight at 4°C then permeabilized with 0.5% Triton X-100 in 1X PBS for 10 min at RT. Cells were incubated with 0.2N HCl for 20 min at RT and washed briefly in 1X PBS. Cells were incubated with 1ug/mL Proteinase K (20mg/mL, ThermoFisher Scientific, cat# AM2546) in Prot K buffer (100mM Tris pH 8.0, 50mM EDTA) for 10 min at 37°C and quenched with 0.2% glycine in 1X PBS for 20 min at RT. Immunostaining was carried out as described above. Cells were imaged in 1X PBS on a Zeiss AxioPhot fluorescence microscope using either 25X/0.80NA or 40X/0.90NA water immersion objective at RT. All fluorescence channels were imaged at non-saturating levels, and settings were kept identical between all samples within replicates used for comparisons or quantifications. For antibodies used refer to “Antibodies and primers” section.

### Combined HSATII RNA-FISH and immunofluorescence

Cells were cultured in 35mm dishes or grown in 4-or 8-well chamber slides. Immunofluorescence was performed as stated in “Immunofluorescence” section with modifications. After secondary staining cells were re-fixed in 4% paraformaldehyde for 7 min at RT. Cells were then washed in hybridization wash buffer (2X SSC, 50% Formamide) for 10 min at RT. Locked nucleic acid, FITC-conjugated HSATII probes were purchased from QIAGEN and are based on the sequence used in previous publications (Hall et al., 2017). Probe: 5′ FAM-ATTCCATTCAGATTCCATTCGATC detects the reverse HSATII transcript. HSATII probes were diluted to 5.0 pmol/mL in whole chromosome painting buffer (50% formamide (Sigma-Aldrich), 2X SSC (Invitrogen), 10% dextran sulfate) and hybridized overnight at 37°C. Cells were washed with 15% formamide/2X SSC for 20 min at 37°C, 2X SSC for 20 min at 37°C and 2X SSC for 5 min at RT. Cells were washed with 1X PBS and nuclei were stained using DAPI for 10 min at RT. Cells in 35mm dishes were imaged in 1X PBS on a Zeiss AxioPhot fluorescence microscope using either 25X/0.80NA or 40X/0.90NA water immersion objective at RT. Cells in chamber slides were briefly air-dried and mounted using ProLong Glass Antifade Mounting with NucBlue (ThermoFisher Scientific). Cells were imaged with a Zeiss Axio Imager Z2 upright microscope as part of a TissueFAXS system (TissueGnostics, Vienna, Austria, software version 7.1.133) using a 40x/0.75 NA Zeiss EC Plan-NEOFLUAR air objective with an ORCA-Flash 4.0 monochrome sCMOS camera. All fluorescence channels were imaged at non-saturating levels, and settings were kept identical between all samples within replicates used for comparisons or quantifications. For antibodies used refer to “Antibodies and primers” section.

### Microscope image acquisition and analysis

Widefield imaging was performed with a Nikon Eclipse Ti fluorescence microscope using a Photometrics Prime BSI Express sCMOS monochrome camera or a Nikon DS-Fi3 color camera using a 20x/0.45NA Air S Plan Fluor ELWD, PH1 ADM objective or a 40x/0.60NA Air S Plan Fluor ELWD, PH2 ADM objective at RT. NIS Elements 5.42.01 acquisition software was used for image acquisition and analysis. Additional widefield imaging was performed using a Zeiss AxioPhot fluorescence microscope with a AxioCam MRm camera (Zeiss) using either 25x/0.80NA or 40x/0.90NA water immersion/ oil objective at RT. AxioVision SE64 Rel. 4.9.1 acquisition software was used for image acquisition and analysis. ImageJ analysis software was used for further analysis including plot profile, mean fluorescence intensity measurements, and co-localization analysis. Proportion of signal overlap between two channels was calculated by segmenting individual nuclei and segmenting foci to be analyzed by thresholding signal and generating ROI from nuclei or foci, then overlap of signal was generated from both channel ROIs. Integrated density measurements were performed on all measured ROIs and analysis of proportion overlap between channels was performed. Protein signal intensity was measured by segmenting individual nuclei or segmenting foci of interest in one channel to be analyzed by thresholding signal and generating ROI. Then signal intensity from second channel was measured in nuclear ROI or foci ROI.

### RNA dot blot

Total RNA was harvested with the NucleoSpin RNA Kit (Takara) according to the manufacturer’s protocol. RNA quality was verified by NanoDrop 2000 (ThermoFisher Scientific). RNA was treated with DNaseI, Amplification Grade (Invitrogen). RNA dot blot was performed as previously described (Shen et al., 2017) with minor modifications. Briefly, RNA was denatured at 95°C for 5 minutes and chilled on ice immediately after denaturation. 2 μL of total RNA was dropped onto an Amersham Hybond-N+ membrane (GE Healthcare, cat# RPN203B), then the membrane was briefly air dried and crosslinked using Stratalinker 2400 UV Crosslinker twice using Autocrosslink mode. Membrane was washed in 10 mL of wash buffer (0.02% Tween-20, 1X PBS) at RT (2x). Membrane was probed for total RNA using methylene blue (Statlab medical products) prior to blocking. Membrane was blocked in 10 mL of blocking buffer (0.02% Tween-20, 5% non-fat milk, 1X PBS) for 1 hour at RT with gentle shaking. Membrane was incubated with anti-m5C antibody in 10 mL of blocking buffer overnight at 4°C. Membrane was washed in 10 mL wash buffer for 15 minutes at RT (2x). Membrane was incubated with HRP-conjugated secondary antibody for 1 hour at RT. Membrane was washed in 10 mL wash buffer for 10 minutes (4x). Membrane was incubated with 5 mL with either SuperSignal West Pico PLUS Substrate (ThermoFisher Scientific) or SuperSignal West Femto Substrate (ThermoFisher Scientific) on a ChemiDoc MP imaging system (BioRad) using ImageLab 6.1 software (BioRad). Blots were analyzed using ImageLab 6.0 software (BioRad) and ImageJ for quantification of signal intensities. For antibodies used refer to “Antibodies and primers” section.

### Gapmer-mediated knockdown

Gapmers were transfected into iDUX4 cells after doxycycline induction using Lipofectamine RNAiMAX reagent (ThermoFisher Scientific) following the manufacturer’s instructions. Gapmers were ordered from QIAGEN. Sequences of gapmers (a ‘+’ indicates a locked nucleic acid modification in the following base): Control_gfp: +g+a+g+aAAGTGTGACA+a+g+t+g, HSATII_F: +a+t+g+gAATCGAATGGA+a+t+c+a, and HSATII_R: +c+a+t+tCGATGATTCC+a+t+t+c.

### Western blot

Cells were harvested in 100μL cold RIPA buffer (150mM NaCl, 1% NP-40, 0.5% sodium deoxycholate, 0.1% SDS, 25mM Tris-HCl pH 7.4) supplemented with Pierce Protease Inhibitors EDTA-free (PIA32955) and Pierce Phosphatase Inhibitors (PIA32957). Cells were scraped and cell lysate was incubated on ice for 15 min. Samples were sonicated in a Biorupter (Diagenode) for 5 min on low, 30 sec on/ 30 sec off to aid in lysis. Samples were centrifuged at 16,000rcf for 10 min at 4°C and supernatant was quantified using Pierce Protein BCA assay kit (ThermoFisher Scientific). Samples were diluted in 1X NuPAGE LDS buffer (Invitrogen)/ 2.5% β-Mercaptoethanol and boiled at 70°C for 10 min. For SDS-PAGE, proteins were loaded onto a 4-12% NuPAGE Bis-Tris gel (Invitrogen) and run at 120V for 1-2 hours in NuPAGE MES SDS Running Buffer or NuPAGE MOPS SDS running buffer (Invitrogen) with 250μL NuPAGE antioxidant (Invitrogen). Proteins were transferred to a 0.2µm or 0.4μM PVDF membrane that was pre-soaked in MeOH for 1min and transferred at 30V or 200mA for 1-1.5hour at 4°C in NuPAGE Transfer Buffer (Invitrogen) containing 20% MeOH. Membranes were blocked in 5% milk/ 1X TBS/ 0.1% Tween or 5% BSA/ 1X TBS/ 0.1% Tween for 1 hour, then incubated with primary antibody in blocking buffer overnight at 4°C with gentle rocking. Blots were washed with 1X TBS/ 0.1% Tween for 15 min at RT (2x) and incubated with HRP-conjugated secondary antibody for 1 hour in blocking buffer. Blots were washed with 1X TBS/ 0.1% Tween for 15 min at RT (2x), and bands were detected via chemiluminescence with either SuperSignal West Pico PLUS Substrate (ThermoFisher Scientific) or SuperSignal West Femto Substrate (ThermoFisher Scientific) on a ChemiDoc MP imaging system (BioRad) using ImageLab 6.1 software (BioRad). Blots were analyzed using ImageLab 6.0 software (BioRad) and ImageJ for quantification of signal intensities. For antibodies used refer to “Antibodies and primers” section.

### Chromatin Isolation by RNA purification (ChIRP) and mass spectroscopy

ChIRP was performed as previously described (Chu et al., 2012) with modifications. 6-10 15 cm plates of cells were used per ChIRP-MS experiment. Cells are cross-linked in 2% paraformaldehyde for 15 minutes at RT, followed by 0.125 M glycine quenching for 5 minutes, scraped and pellet at 650xg for 5 minutes at 4°C. Cells were then lysed in lysis buffer (50 mM Tris-HCl, pH 7.0, 10 mM EDTA, pH 8.0, 1% SDS supplemented with fresh 0.5 mM AEBSF (ThermoFisher Scientific, cat# BP635-500), Pierce Protease Inhibitors EDTA-free (PIA32955) and Pierce Phosphatase Inhibitors (PIA32957) and RNase inhibitor (TaKaRa, cat# 2313A)) for 10 minutes on ice. Samples were sonicated in a Biorupter (Diagenode) for 15 minutes on high, 30 sec on/ 30 sec off pulse intervals for 4-cycles. Samples were centrifuged at 16,000xg for 10 minutes at 4°C and 5% of the lysate was taken out as an input control. Lysates were pre-cleared with beads (Invitrogen Dynabeads M-280 Streptavidin, cat# 11205D) at 37°C for 30 minutes at 800rpm. Hybridization was performed at 37°C for 4 hours shaking at 800rpm using biotin-conjugated probes (HSATII probe: 5′-ATTCCATTCAGATTCCATTCGATC-3BioTEG-3’, Control probe: 5’-GTCCCGTTAGCTCAGGTGGTAGAGCAC-3BioTEG-3’ or 5’-TGCTGATGAAGCAGAACAAC-3BioTEG-3’) in hybridization buffer (750 mM NaCl, 1% SDS, 50 mM Tris-HCl, pH 7.0, 1 mM EDTA, 15% formamide (Sigma-Aldrich) and supplemented with fresh 0.5 mM AEBSF (ThermoFisher Scientific, cat# BP635-500), Pierce Protease Inhibitors EDTA-free (PIA32955) and Pierce Phosphatase Inhibitors (PIA32957) and RNase inhibitor (TaKaRa, cat# 2313A)). Streptavidin magnetic beads (Invitrogen Dynabeads M-280 Streptavidin, cat# 11205D) were added (100 mL per 100 pmole of probes used) and incubated at 37°C for 30 minutes at 800rpm. Beads were washed 3x in pre-warmed (37°C) wash buffer (2X SSC (Invitrogen), 0.5% SDS and supplemented with Pierce Protease Inhibitors EDTA-free (PIA32955) and Pierce Phosphatase Inhibitors (PIA32957) and RNase inhibitor (TaKaRa, cat# 2313A) and 1 mM DTT) for 5 minutes at 37°C at 800rpm. 10% of beads were removed for RNA isolation and 90% of remaining beads were used for protein isolation. RNA was isolated in RNA elution buffer (10 mM EDTA, pH 8.0, 95% formamide (ThermoFisher Scientific)) and incubated at 95°C for 5 minutes at 1,000xg. Proteinase K was added (20mg/mL, ThermoFisher Scientific, cat# AM2546) and incubated at 55°C for 1 hour. RNA was extracted using TRIzol and validation of *HSATII* enrichment was performed using RT-qPCR. Protein was isolated by boiling beads for 30 minutes at 95°C in 2X NuPage LDS sample buffer (ThermoFisher Scientific, cat#NP0007). Protein samples were using in immunoblotting or for LC-MS. For LC-MS, eluted protein samples were electrophoresed into a NuPage 4–12% Bis-Tris gel, excised, and processed by the Fred Hutchinson Cancer Research Center Proteomics Core. Samples were reduced, alkylated, digested with trypsin, desalted, and run on the Orbitrap Eclipse Tribid Mass Spectrometer (ThermoFisher Scientific). Proteomics data were analyzed using Proteome Discoverer 2.5 against a UniProt human database that included common contaminants using Sequest HT and Percolator for scoring. Results were filtered to only include protein identifications from high-confidence peptides with a 1% false discovery rate. Proteins that were identified in all replicates from both independent experiments with at greater than ten unique peptide matches and greater than 1.5 difference between sample and control were analyzed further.

### RNA Interactome using Click Chemistry (RICK)

RICK was performed as previously described (Bao et al., 2018). Briefly, cells were cultured in 15cm plates and 0.1mM 5-ethynyl uridine (ThermoFisher Scientific) was incubated with cells after dox-induction for 16-hrs followed by a washout and then fixed at 48-hrs. After washing 3x with 1X PBS, the plates were placed on ice and irradiated with 0.15 J/cm^2^ UV light at 254 nm (Stratalinker). Cells were then fixed with 90% ethanol for 30 min, washed 3x with 1X PBS, and permeabilized with 0.5% Triton X-100 in 1X PBS for 15 min. Permeabilized cells were incubated with 15 mL of click reaction buffer (0.25M biotin-azide (Click Chemistry Tools, cat# 1265-25), 0.5M CuSO4, 0.25M THPTA, 0.1M sodium L-ascorbate in 1X PBS) for 30 minutes at RT. Reaction was quenched 3x in 0.5% Triton X-100 in 1X PBS with 2mM EDTA for 3 minutes at RT. Cells were then lysed in lysis buffer (20 mM Tris–HCl, pH 7.5, 500 mM LiCl, 1 mM EDTA pH 8.0, 0.5% lithium-dodecylsulfate (LiDS), and 5 mM DTT) supplemented with Pierce Protease Inhibitors EDTA-free (PIA32955) and Pierce Phosphatase Inhibitors (PIA32957) and RNase inhibitor (TaKaRa, cat# 2313A), and harvested by scraping. Samples were sonicated in a Biorupter (Diagenode) for 15 min on low, 30 sec on/ 30 sec off to aid in lysis. Samples were centrifuged at 12,000 rcf for 10 min at 4°C and 5% of the lysate was taken out as an input control. Complexes containing different RNA species, and their associated proteins, were isolated with streptavidin-conjugated magnetic beads (100 μl beads for each plate; Dynabeads MyOne Streptavidin C1, Thermo Fisher Scientific, cat# 65602). After incubation with the lysates overnight under continuous rotation at 4°C, the beads were isolated on a magnetic stand and washed using lysis buffer, buffer 1 (20 mM Tris–HCl, pH 7.5, 500 mM LiCl, 1 mM EDTA pH 8.0, 0.1% LiDS, and 5 mM DTT), buffer 2 (20 mM Tris–HCl pH 7.5, 500 mM LiCl, 1 mM EDTA pH 8.0, and 5 mM DTT), and buffer 3 (20 mM Tris–HCl pH 7.5, 200 mM LiCl, 1 mM EDTA pH 8.0, and 5 mM DTT) for two times each under rotation (for 10 minutes at 4°C). At last wash step beads were split to isolate RNA or proteins. Proteins were extracted from the captured complexes using RNase A/T1 mix (ThermoFisher Scientific, cat# EN0551) and Ambion RNase III (ThermoFisher Scientific, cat# AM2290) at 37°C for 1 hour at 1,000xg, then boiled for 10 minutes 95°C. RNA was isolated in RNA elution buffer (10 mM EDTA, pH 8.0, 95% formamide (ThermoFisher Scientific)) and incubated at 95°C for 5 minutes at 1,000xg. Proteinase K was added (20mg/mL, ThermoFisher Scientific, cat# AM2546) and incubated at 55°C for 1 hour. RNA was extracted using TRIzol.

### RICK liquid chromatography mass spectroscopy (LC-MS)

For LC-MS, eluted protein samples were electrophoresed into a NuPage 4–12% Bis-Tris gel, excised, and processed by the Fred Hutchinson Cancer Research Center Proteomics Core. Samples were reduced, alkylated, digested with trypsin, desalted, and run on the Orbitrap Fusion Mass Spectrometer (ThermoFisher Scientific). Proteomics data were analyzed using Proteome Discoverer 2.4 against a UniProt human database that included common contaminants using Sequest HT and Percolator for scoring. Results were filtered to only include protein identifications from high-confidence peptides with a 1% false discovery rate. Proteins that were identified in all replicates from both independent experiments with at least two unique peptide matches and greater than 1.5 difference between sample and control were analyzed further.

### HSATII KD RNA sequencing and data analysis

Total RNA was harvested with the NucleoSpin RNA Kit (Takara) according to the manufacturer’s protocol. RNA was treated with DNaseI, Amplification Grade (Invitrogen). RNA quality was verified by NanoDrop 2000 (ThermoFisher Scientific) and Tapestation (Agilent). TruSeq stranded mRNA library preparation was done by the Fred Hutchinson Cancer Research Center Genomics core. Libraries were sequenced using 100 bp single-end sequencing on the Illumina NextSeq P2 platform by the Fred Hutchinson Cancer Center Genomics core facility. RNA-seq analysis was done in a similar fashion as the EU labeled RNA-seq above with volcano plots produced using EnhancedVolcano (Blighe et al., 2024). Differential isoform usage and splicing analysis was done using IsoformSwitchAnalyzeR (version 2.4.0) (Vitting-Seerup and Sandelin, 2017; Vitting-Seerup and Sandelin, 2019) and Kallisto (version 0.48.0) (Bray et al., 2016) along with Gencode annotation release R45. Settings for Kallisto pseudo-alignment were “-*-l 200 –-s 200 –-plaintext –-bootstrap-samples=10 – –pseudobam –-genomebam –gtf gencode.v45.primary_assembly.annotation.gtf”.* Gene set enrichment analysis was analyzed using Enrichr (Chen et al., 2013; Kuleshov et al., 2016; Xie et al., 2021).

### Quantitative RT-PCR

RNA was harvested with the NucleoSpin RNA Kit (Takara) according to the manufacturer’s protocol. RNA quality was verified by NanoDrop 2000 (ThermoFisher Scientific). RNA was treated with DNaseI, Amplification Grade (Invitrogen). Reverse transcription was performed using the Superscript IV First-Strand Synthesis System. For 20uL reaction: 200-1000ng total RNA, 1μL 10mM dNTPs, 1μL 10mM random hexamers, 4μL 5X SSIV Buffer, 1μL 100mM DTT, 1μL RNaseOUT (Invitrogen), and 1μL SSIV RT enzyme. Thermal cycling conditions for reverse transcription were as follows: 50°C for 40 min, 55°C for 30 min, and 80°C for 10 min. Complementary DNA (cDNA) was treated with 1μL of RNaseH and incubated at 37°C for 20 min, then diluted 1:5 with RNase-free H_2_O. Quantitative real-time PCR (qPCR) was performed on the QuantStudio™ 7 Flex Real-Time PCR System in a 10μL reaction: 2μL cDNA, 5μL 2X iTaq Universal SYBR Green Supermix, 0.3μL 10μM forward and reverse primer, and 2.4μL H_2_O. qPCR primers were synthesized by Integrated DNA Technologies (IDT) and are listed in “Antibodies and primers” section. Thermal cycling conditions for qPCR were as follows: 50°C for 2 min and 95°C for 10 min; 40 cycles of 95°C for 15 sec and 60°C for 60 sec. For each biological replicate, qPCR reactions were run in technical triplicates, including –RT controls. Median CT values of the technical triplicates were used for analysis. Gene expression was normalized to the housekeeping gene RPL27 (ribosomal protein L27). For primers used refer to “Antibodies and primers” section.

### siRNA knockdown

siRNAs were transfected into iDUX4 cells 24-to 72-hours prior to doxycycline induction using Lipofectamine RNAiMAX reagent (ThermoFisher Scientific) following the manufacturer’s instructions. ON-TARGETplus SMARTpool siRNAs were ordered from Horizon Discovery/ Dharmacon Reagents. siRNAs used: Human YBX-1 (Catalog ID L-010213-00-0005), Human NSUN2 (L-018217-01-0005), Human KIAA1429 (VIRMA; L-019278-02-0005) and control (D-001810-10).

### Statistics

All statistical analyses were performed in Prism (GraphPad). Bars represent mean ± SD. At least three experimental replicates were performed unless otherwise stated. Statistical measures are described in the figure legends.

### Antibodies and primers

*Primary antibodies:* SC35 (Abcam Cat# ab204916, RRID:AB_2909393), TDP-43 (Proteintech Cat# 10782-2-AP, RRID:AB_615042), MeCP2 (Abcam Cat#ab137357), NSUN2 (Proteintech Cat# 20854-1-AP, RRID:AB_10693629), NSUN2 (Cell Signaling Techonology Cat#52901), MVP (Proteintech Cat# 16478-1-AP, RRID:AB_2147597), YBX-1 (Abcam Cat# ab76149, RRID:AB_2219276), NPM1 (Proteintech Cat# 60096-1-Ig, RRID:AB_2155162), FBL (Proteintech Cat# 16021-1-AP, RRID:AB_2105788), m^5^C (Cell Signaling Technology Cat# 28692, RRID:AB_2798962), K1 (SCICONS Cat# 10020200, RRID:AB_2756865), GAPDH (GeneTex Cat# GTX28245, RRID:AB_370675), eIF4A3 (Abcam Cat#ab180573), H3.X/Y (Active Motif Cat# 61161, RRID:AB_2793533), Beta Tubulin (Proteintech Cat# 10094-1-AP, RRID:AB_2210695), VIRMA (Proteintech Cat# 25712-1-AP, RRID:AB_2880204), WTAP (Cell Signaling Technology Cat# 56501, RRID:AB_2799512), YTHDC1 (Cell Signaling Technology Cat# 77422, RRID:AB_2799899), m^6^A (Synaptic Systems Cat# 202 003, RRID:AB_2279214), DUX4 (E14-3) (Geng et al., 2012), eIF4ENIF1 (Thermo Fisher Scientific Cat# PA5-115169, RRID:AB_2899805), TRIM21 (Thermo Fisher Scientific Cat# PA5-22294, RRID:AB_11153206), Goat Anti-Rabbit IgG (H+L) Superclonal Recombinant Secondary Antibody, HRP (Thermo Fisher Scientific Cat# A27036, RRID:AB_2536099), Goat Anti-Mouse IgG (H+L) Superclonal Recombinant Secondary Antibody, HRP (Thermo Fisher Scientific Cat# A28177, RRID:AB_2536163), Goat anti-Rat IgG (H+L) Secondary Antibody, HRP (Thermo Fisher Scientific Cat# 31470, RRID:AB_228356), Rhodamine (TRITC)-AffiniPure Donkey Anti-Rabbit IgG (H+L) (Jackson ImmunoResearch Labs Cat# 711-025-152, RRID:AB_2340588), Rhodamine (TRITC)-AffiniPure Donkey Anti-Mouse IgG (H+L) (Jackson ImmunoResearch Labs Cat# 715-025-151, RRID:AB_2340767), Fluorescein (FITC)-AffiniPure Donkey Anti-Mouse IgG (H+L) (Jackson ImmunoResearch Labs Cat# 715-095-151, RRID:AB_2335588), Fluorescein (FITC)-AffiniPure Donkey Anti-Rabbit IgG (H+L) (Jackson ImmunoResearch Labs Cat# 711-095-152, RRID:AB_2315776), Streptavidin, Alexa Fluor 555 conjugate antibody (ThermoFisher Scientific Cat# S21381), Donkey anti-Rabbit IgG (H+L) Highly Cross-Adsorbed Secondary Antibody, Alexa Fluor 488 (Thermo Fisher Scientific Cat# A-21206, RRID:AB_2535792), Donkey anti-Mouse IgG (H+L) Highly Cross-Adsorbed Secondary Antibody, Alexa Fluor 488 (Thermo Fisher Scientific Cat# A-21202, RRID:AB_141607), Donkey anti-Rabbit IgG (H+L) Highly Cross-Adsorbed Secondary Antibody, Alexa Fluor 555 (Thermo Fisher Scientific Cat# A-31572, RRID:AB_162543), Donkey anti-Mouse IgG (H+L) Highly Cross-Adsorbed Secondary Antibody, Alexa Fluor 555 (Thermo Fisher Scientific Cat# A-31570, RRID:AB_2536180).

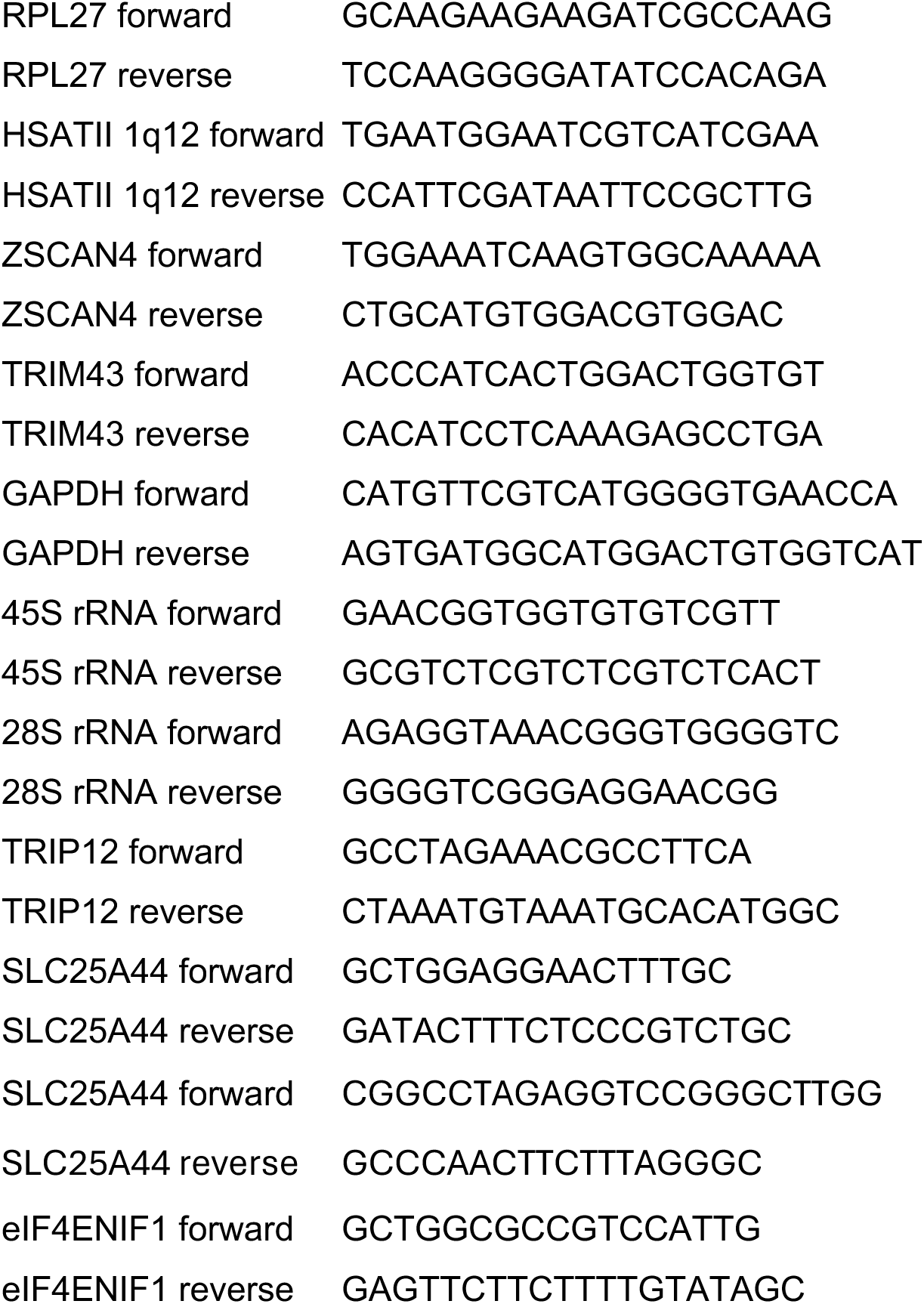
*Primers:* 5’-NNN-3’.

## SUPPLEMENTAL FIGURE LEGENDS

**Supplemental Figure 1.**
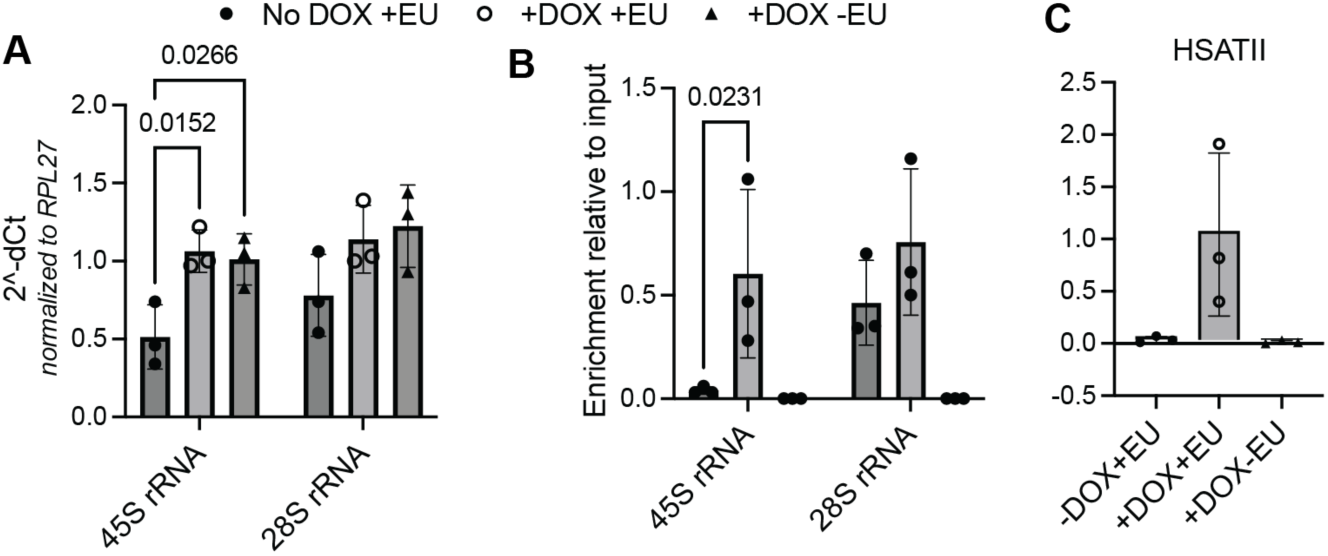
Identification of DUX4-induced EU-labeled RNA aggregates. (**A**) Relative expression of either 45S precursor ribosomal RNA (rRNA) or mature 28S rRNA from isolated EU-RNA from –DOX or +DOX iDUX4 cells that were incubated with EU (+EU) or without EU (-EU) for 16-hours and then harvested at 48-hours. Dots represent individual experimental replicates. N=3. Data represent means ± SD. Statistical differences between groups were analyzed employing one-way ANOVA Dunnett’s multiple comparison test. (**B**) Relative enrichment of either 45S precursor rRNA or mature 28S rRNA in isolated EU-RNA fraction compared to input from –DOX or +DOX iDUX4 cells that were incubated with EU (+EU) or without EU (-EU) for 16-hours and then harvested at 48-hours. Dots represent individual experimental replicates. N=3. Data represent means ± SD. Statistical differences between groups were analyzed employing one-way ANOVA Dunnett’s multiple comparison test. (**C**) Enrichment of *HSATII* RNA from isolated EU-RNA from –DOX or +DOX iDUX4 cells that were incubated with EU (+EU) or without EU (-EU) for 16-hours and then harvested at 48-hours. Dots represent individual experimental replicates. N=3. Data represent means ± SD.

**Supplemental Figure 2.**
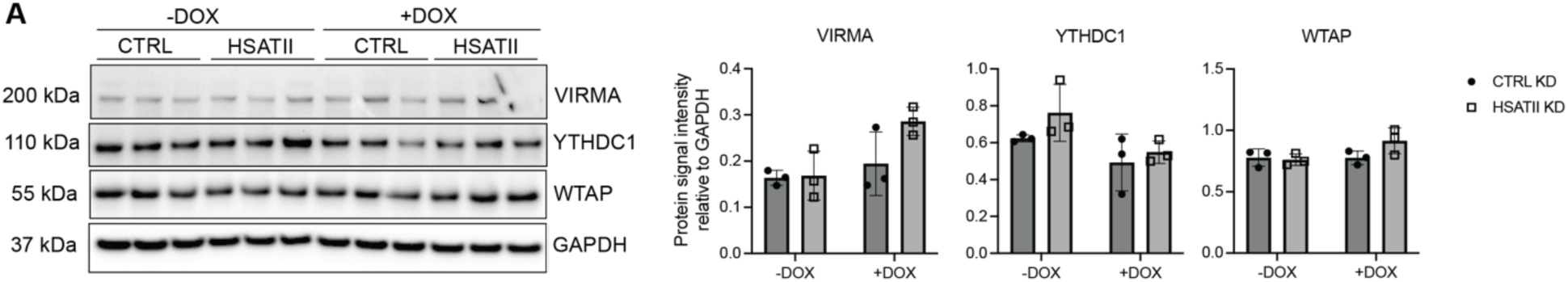
m^6^A-related protein levels in DUX4-expressing cells. (**A**) Immunoblot of whole cell lysate from –DOX or +DOX iDUX4 cells with CTRL KD or HSATII KD and harvested at 24-hours post-induction. Blot was probed for VIRMA, YTHDC1 and WTAP total protein levels. GAPDH was used as loading control. Immunoblot shows three biological replicates, which is representative of two experimental replicates performed. Quantification of relative protein levels are indicated adjacent to blot.

**Supplemental Figure 3.**
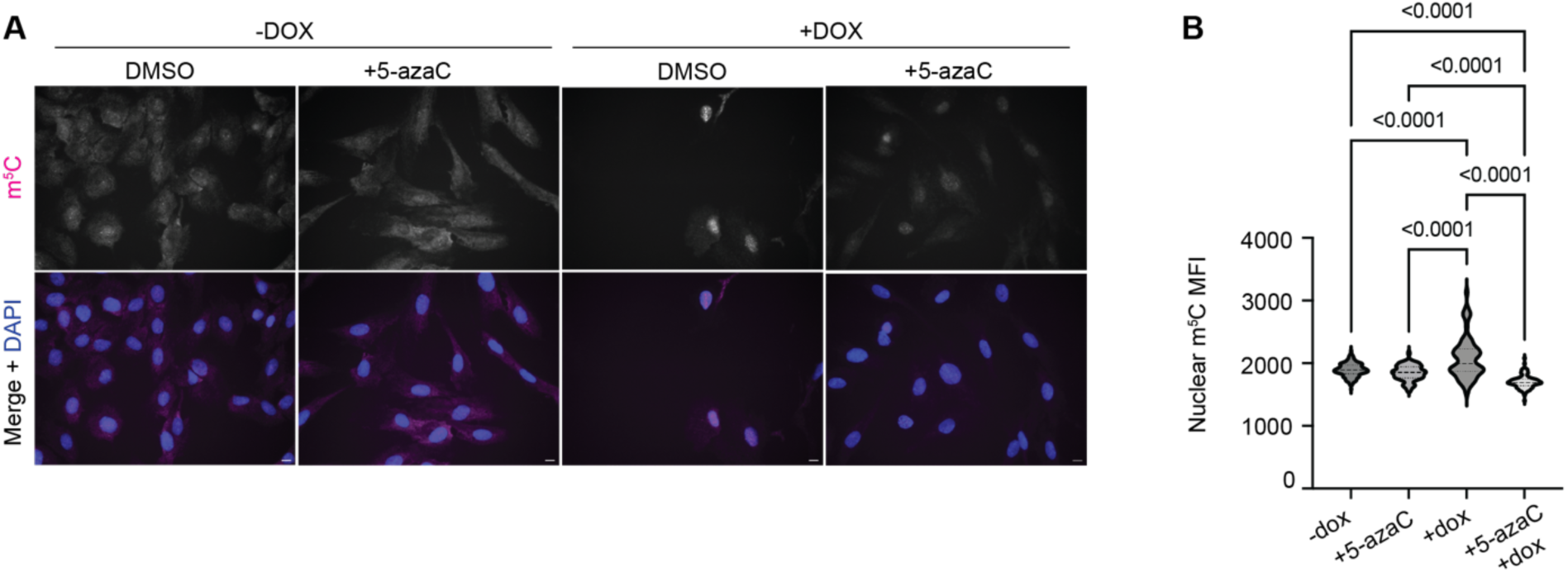
Treatment with 5-azacytidine in iDUX4 cells. (**A**) Immunofluorescence of m^5^C (magenta) in –DOX or +DOX iDUX4 cells either pre-treated with DMSO or 5μM 5-azacytidine for 24-hours, then dox-induced and fixed 24 hours post-induction. Scale bar = 20μm. Images are representative of two independent experimental replicates performed. (**B**) Nuclear m^5^C mean intensity in –DOX or +DOX iDUX4 cells either pre-treated with DMSO or 5μM 5-azacytidine for 24-hours, then dox-induced and fixed 24 hours post-induction. N ≥ 70 nuclei. Data are representative of two independent experimental replicates. Statistical differences between –DOX (control) and +DOX iDUX4 cells were analyzed employing one-way ANOVA Tukey’s multiple comparison test.

**Supplemental Figure 4.**
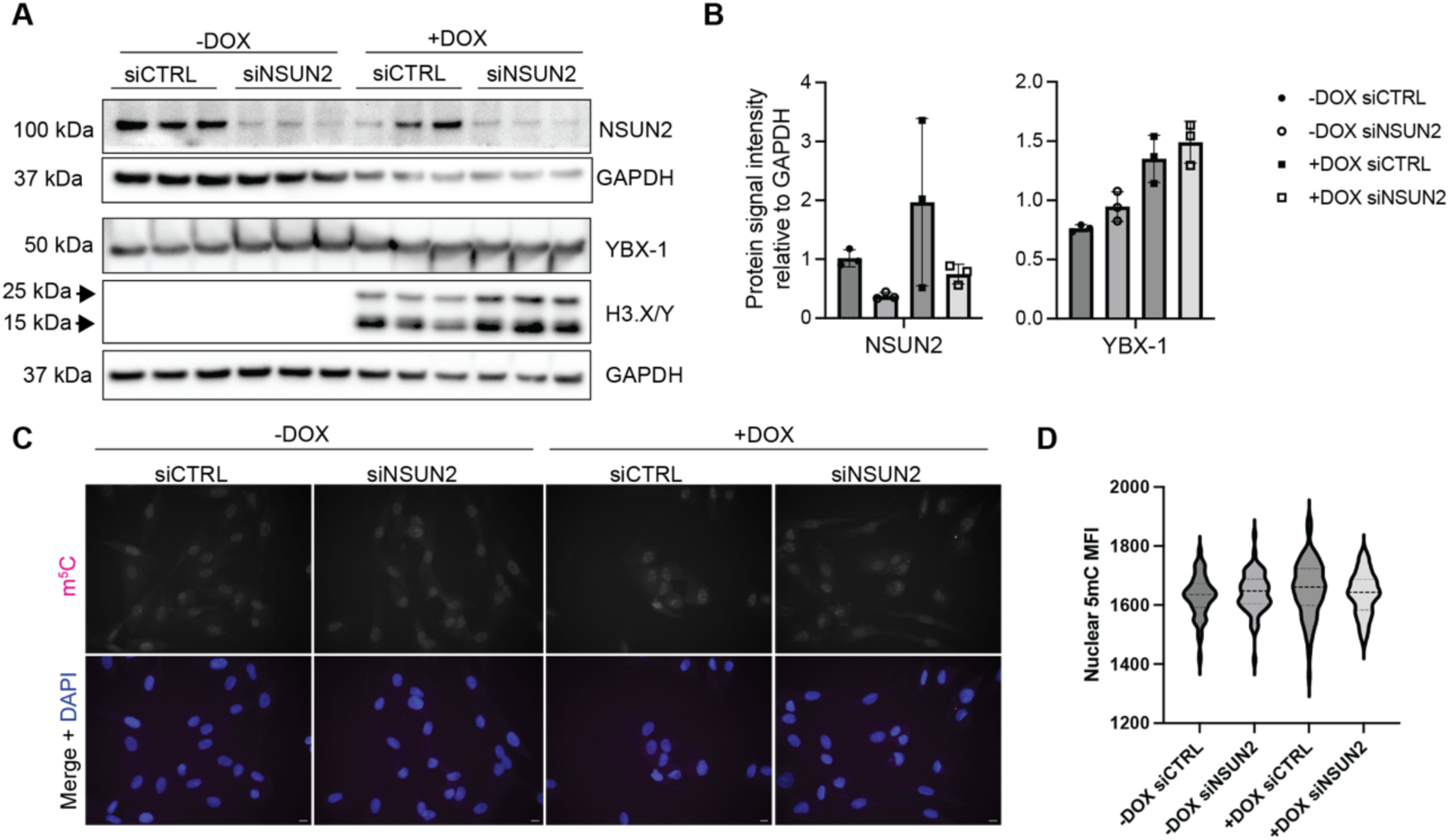
NSUN2 depletion does not impact YBX-1 protein levels or global m^5^C nuclear signal. (**A**) Immunoblot of NSUN2, YBX-1 and H3.X/Y protein levels from whole cell lysate from –DOX or +DOX iDUX4 cells with control depletion (siCTRL) or NSUN2 depletion (siNSUN2). GAPDH was used as loading control. Immunoblot shows three biological replicates, which is representative of two experimental replicates performed. (**B**) Protein levels NSUN2 and YBX-1 relative to GAPDH loading control of immunoblot shown in (A). Dots indicate biological replicate. (**C**) Immunofluorescence of m^5^C (magenta) in –DOX or +DOX iDUX4 cells either pre-treated with siRNAs targeting control sequences (siCTRL) or NSUN2 (siNSUN2) for 24-hours, then dox-induced and fixed 24 hours post-induction. Scale bar = 20μm. Images are representative of two independent experimental replicates performed. (**D**) Nuclear m^5^C mean intensity in –DOX or +DOX iDUX4 cells either pre-treated with siRNAs targeting control sequences (siCTRL) or NSUN2 (siNSUN2) for 24-hours, then dox-induced and fixed 24 hours post-induction. N ≥ 100 nuclei. Data are representative of two independent experimental replicates.

**Supplemental Figure 5.**
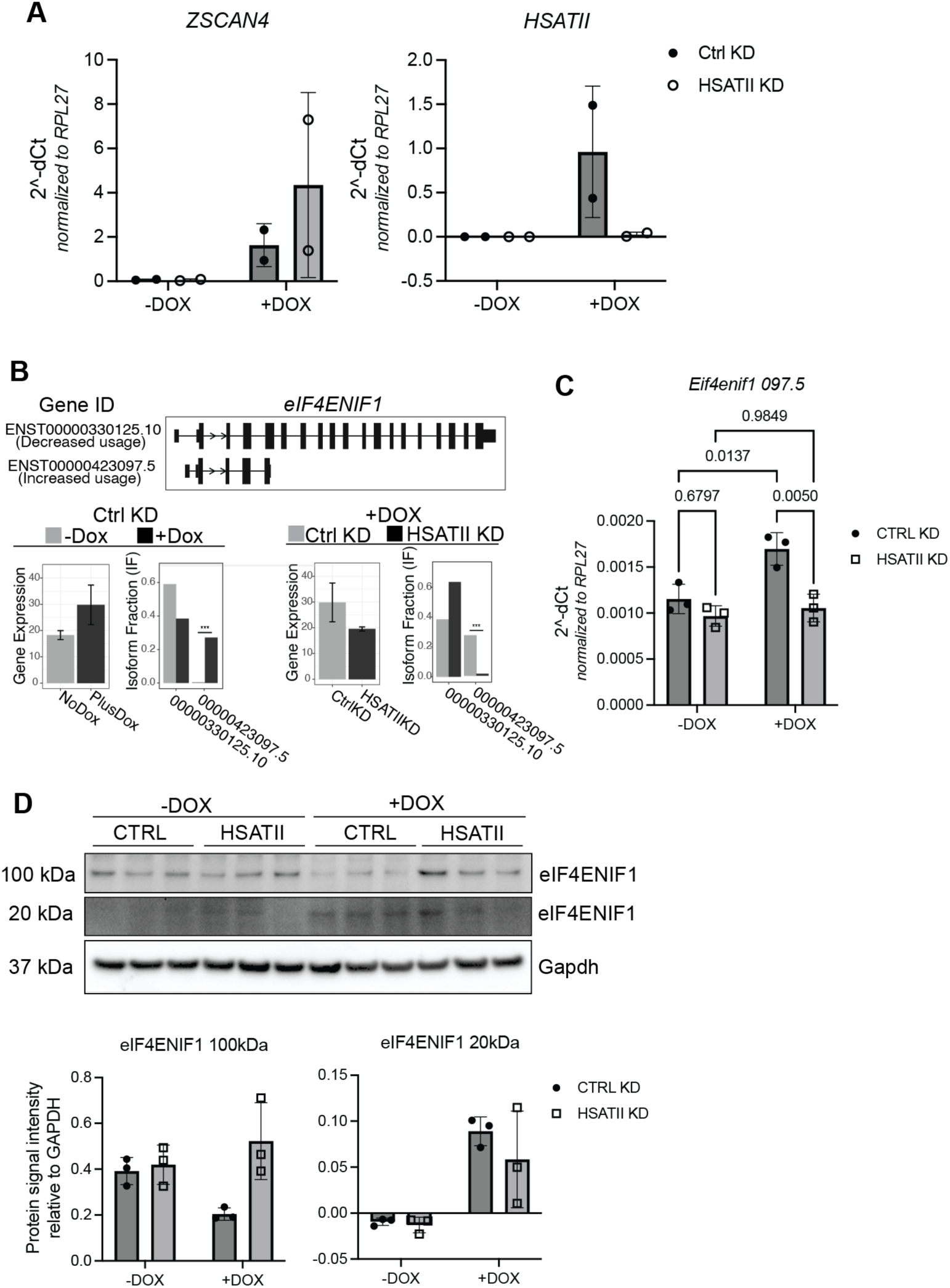
Identification of RNA splicing changes mediated by HSATII-RNP. (**A**) Relative expression of *Zscan4* (DUX4-target and negative control for HSATII-specific depletion) and *HSATII* from isolated total RNA from –DOX or +DOX iDUX4 cells treated with control knockdown (Ctrl KD; black filled-in circle) or HSATII-specific depletion (HSATII KD; black outlined circle). Samples were used for RNA-sequencing experiments. Dots indicate biological duplicates. (**B**) Top panel: *eIF4ENIF1* transcript isoforms and their associated Ensembl gene ID; bottom left panel: changes in *eIF4ENIF1* transcript isoform usage in –DOX versus +DOX iDUX4 cells with CTRL KD. (Left) Gene expression of *eIF4ENIF1*. TPM: transcripts per million. (Right) Isoform usage; bottom right panel: changes in *eIF4ENIF1* transcript isoform usage in +DOX iDUX4 cells with CTRL KD or HSATII KD. (Left) Gene expression of *eIF4ENIF1*. TPM: transcripts per million. (Right) Isoform usage. (**C**) Expression of *eIF4ENIF1* transcript isoform (ensemble gene ID# 00000423097.5) in –DOX or +DOX iDUX4 cells with (HSATII KD) or without (CTRL KD) HSATII depletion. N= 3. Data represent means ± SD. Statistical differences between groups were analyzed employing 2-way ANOVA Tukey’s multiple comparison test. (**D**) Protein expression of eIF4ENIF1 in –DOX or +DOX iDUX4 cells with (HSATII KD) or without (CTRL KD) HSATII depletion. N= 3. Immunoblot shows two protein isoforms that correspond to the predicted translated products of their respective RNA isoforms. Quantitation of protein signal intensity for both protein products are shown below immunoblot.

